# EpicTope: predicting and validating non-disruptive epitope tagging sites

**DOI:** 10.1101/2024.03.03.583232

**Authors:** Joseph Zinski, Parnal Joshi, Henri Chung, James Preston, Ari Geller, Finn Warrick, Brian D. Berg, K. Tobin, Greg Glova, Fang Liu, Zhitao Ming, Maura McGrail, Darius Balciunas, Iddo Friedberg, Mary C. Mullins

**Author notes:** Co-corresponding authors @.

## Abstract

Epitope tagging is a valuable technique enabling the *in vivo* identification, tracking, and purification of proteins. We developed a tool, EpicTope, to facilitate this method by identifying amino acid positions most suitable for epitope insertion. Our method uses a scoring function that considers protein sequence secondary and tertiary structural features, solvent accessibility, and disordered binding regions to determine locations least disruptive to the protein’s function. We validated our approach on the zebrafish Smad5 and Hdac1 proteins. We show that multiple predicted internally tagged Smad5 proteins rescue zebrafish *smad5* mutant embryos, while the N- and C-terminal tagged variants do not, as predicted. Similarly, we found that optimally-predicted internal and C-terminal Hdac1 tags rescued *hdac1* mutant embryos, while a less-optimal N-terminal tag did not. We further show that these functionally tagged Smad5 and Hdac1 proteins are accessible to antibodies in wholemount zebrafish embryo immunohistochemistry, by western blot, and by immunoprecipitation from embryo extracts. Our work demonstrates that EpicTope is an accessible and effective tool for designing epitope tag insertion sites.

## Introduction

Scientists rely on antibodies with high sensitivity to specifically recognize proteins for a multitude of functions, including visualizing subcellular localization, assessing expression levels, mapping cellular trafficking, and identifying physical interactions. Unfortunately, high quality antibodies against the majority of the vertebrate proteome are not commercially available. Similarly, endeavors by labs to generate custom antibodies is a time- and resource-consuming process that often does not succeed, leading to a significant loss of research resources. A more reliable option is to engineer an epitope-tagged version of the protein; however, even this approach has drawbacks. Simple N- and C-terminal tags can often fail when the ends are functionally or structurally important (Zordan et al., 2015). Conversely, generating libraries of internal tags via transposon mutagenesis is effective but laborious (Zordan et al., 2015). There is a pressing need for accessible, accurate, validated algorithms to predict sites for epitope tagging that will not disrupt protein function.

There are a bevy of available online resources that could potentially be leveraged to help predict suitable epitope-tag locations in proteins of interest including sequence conservation (MUSCLE), predicted secondary structure (AlphaFold2), predicted solvent accessibility, and predicted disordered regions (IUPred2). A few studies by our lab and others have shown promising predictive results by using sequence conservation (Burg et al., 2016, Gibb et al., 2018) and a combination of sequence conservation and surface accessibility (Oesterle et al., 2017). These and other genetic features have been successfully used to build predictive models for frameshift mutations in humans that cause disease (Folkman et al., 2015; Pagel et al., 2019).

Here, we provide a computational tool, EpicTope, which integrates predictions of tertiary structure, secondary structure, solvent accessibility, disordered binding regions, and evolutionary conservation to predict optimal sites for epitope tagging without disrupting function. To test its efficacy, we used EpicTope to predict the best sites to epitope tag the Smad5 and Hdac1 proteins in zebrafish. Smad5 is an essential downstream transcription factor of the TGF-ꞵ/BMP signaling pathway, first required during gastrulation in zebrafish to specify ventral-lateral axial cell fates. It is highly conserved evolutionarily, with 90% amino acid identity among multiple vertebrates (Hata & Chen, 2016), indicating high conservation pressure and underscoring how difficult it could be to engineer a functional epitope-tagged protein. We found that EpicTope-predicted functional V5-tags at two internal sites of zebrafish Smad5, fully rescued zebrafish *smad5* mutant embryos, while N- and C-terminally tagged Smad5 sites, predicted to have reduced functionality, did not rescue. We further tested EpicTope-predicted epitope tags of Histone deacetylase 1 (Hdac1), an essential protein important in epigenetics and transcriptional regulation. Similarly, we found that V5 tags inserted at the two best predicted sites, an internal and C-terminal site, could functionally rescue *hdac1* mutant embryos. On the other hand, an N-terminal tag with lower predicted functionality did not rescue. We then showed that the functionally-tagged Smad5 and Hdac1 proteins were nuclearly localized in the embryo as expected, detectable on western blots, and by immunoprecipitation from embryo extracts, showing the versatility of the epitope tags. Together, our work provides an accessible tool that can be adapted for use with a wide-range of organisms for predicting optimal epitope-tag sites.

## Results and Discussion

### EpicTope prediction for Smad5

We first applied the EpicTope tagging method (Fig 1) to zebrafish Smad5 (Fig 2). The Smad5 amino acid sequence is unusually highly conserved across vertebrates (Fig S1), with 90% identical residues, with less conserved regions between positions 179-193 and 243-250 (Fig 2A). AlphaFold2 predicts a disordered structure from 166-258 at high confidence, with structured regions flanking this area (Fig 2B). Using the multiple sequence alignment, AlphaFold2 structure, and predictions from IUPred2A, we calculated Shannon entropy, secondary structure, relative solvent accessibility (RSA), and disordered binding region (DBR) feature scores for each position (Fig 2C). While the 166-258 region scores well for both secondary structure and relative solvent accessibility (i.e. no secondary structure predicted and the region is mostly solvent accessible), there is a predicted disordered binding region (and therefore decreased suitability for tagging) in a narrow region between 221-235 (Fig 2B, green line).

**Figure 1.**
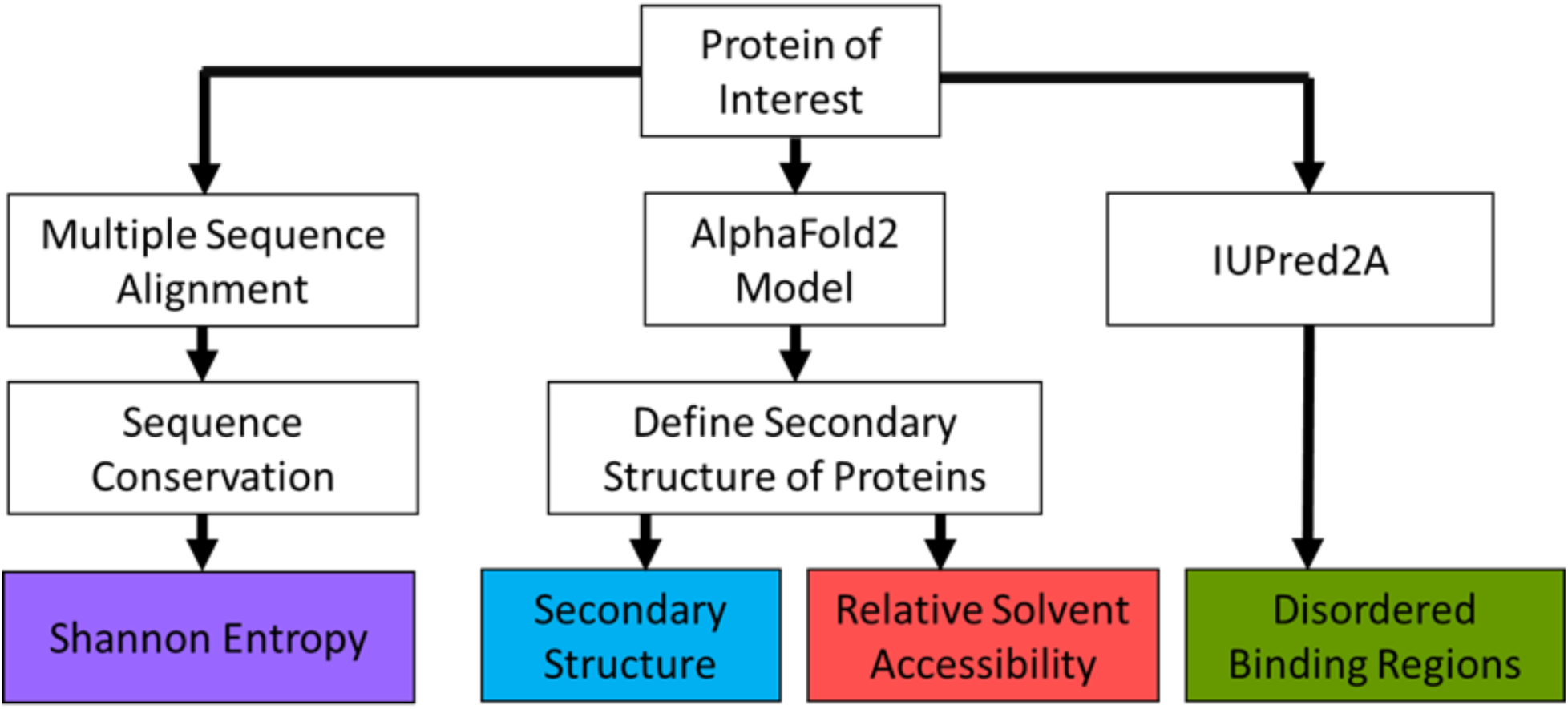
Features used to predict epitope tagging positions. We first identify a protein of interest by its corresponding Uniprot entry. We then retrieve the amino acid sequence and AlphaFold2 predicted Protein Data Bank (PDB) structure. The PDB structure is then used in the Dictionary of Secondary Structure of Proteins (DSSP) program to determine its secondary structures and relative soluble surface area along its sequence. We use a multiple sequence alignment of the amino acid sequence with homologs in seven other species to determine sequence conservation. The UniprotID is used to retrieve the ANCHOR2 score, a measure of disordered protein binding regions, from the IUPRED2A web server.

**Figure 2.**
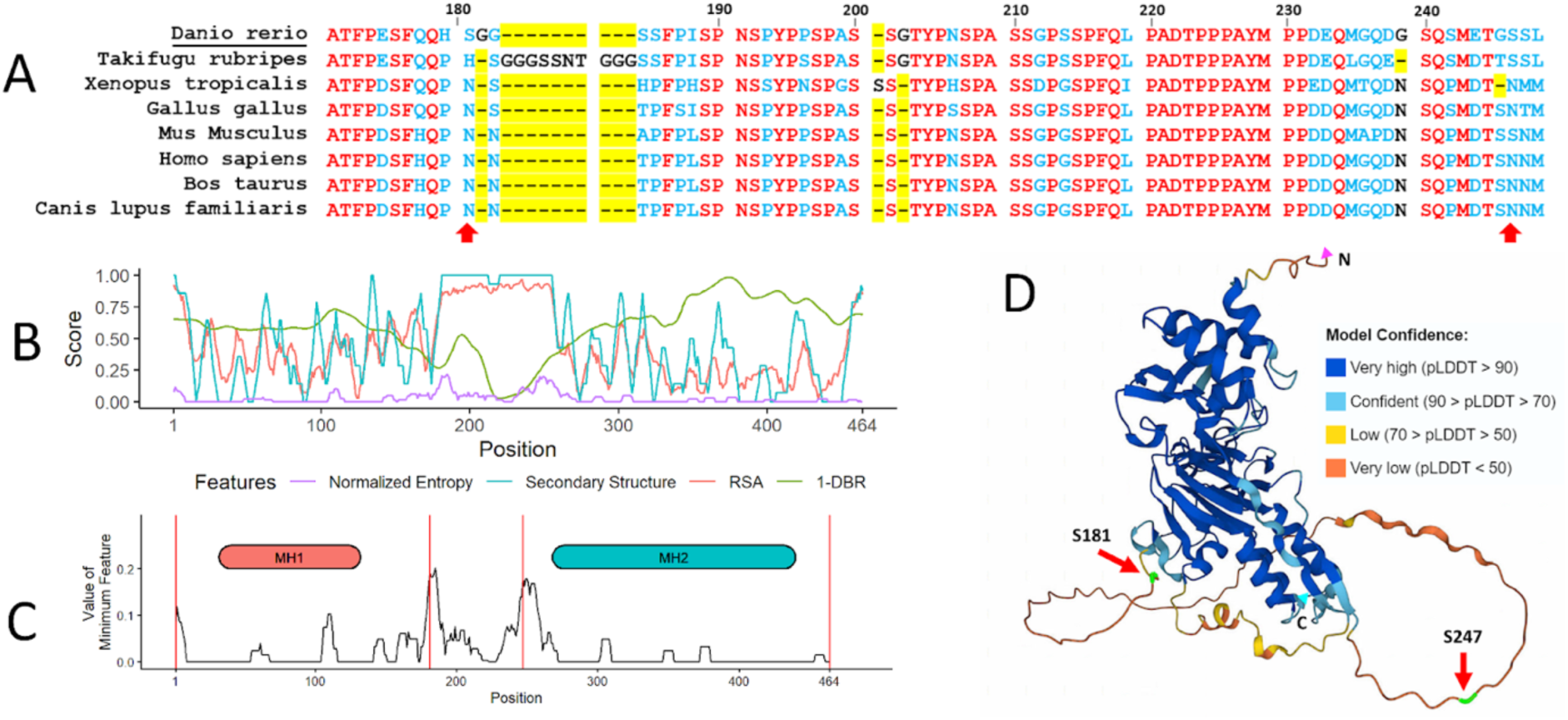
Smad5 EpicTope features and predictions. We calculated EpicTope predictions for Smad5, UniprotID: Q9W7E7. A) Multiple sequence alignment for Smad5 positions 170-250. Amino acids identical between all species are shown in red, differences are in blue. Amino acid insertions or deletions (length variation) are highlighted in yellow. Arrows indicate position of tag inserts. Position indices are labeled in reference to zebrafish *Danio rerio*. B) Shannon entropy, secondary structure, relative solvent accessibility (RSA) and disordered binding region (DBR) features used in EpicTope prediction. Features are normalized to a 0-1 scale and weighted equally. A higher feature score indicates suitability for tagging. C). We plot the minimum score among the four feature scores (*Ei*, see Eq. 5) at each amino acid position. We then select the position(s) with the highest values (181 and 247 here) as the least disruptive tag insertion sites. For each feature, values were averaged along a sliding window of 7 amino acids, except at N- and C-termini where the four terminal amino acids were used. Positions 181, 247, and N-and C-termini are highlighted with red vertical lines. D) Smad5 AlphaFold2 predicted structure; red arrows indicate EpicTope-predicted tagging positions, highlighted in green. Disordered regions are characterized by a lower pLDDT value, a measure of model confidence. C and N termini are labeled with blue and pink triangles.

We score tagging suitability in two ways. First, we identify the minimum score among the four features for each amino acid position, and then we identify the highest minimum scoring positions along the sequence. Raw scores of the minimum scoring feature at S181 and S247 are 0.23 and 0.2, respectively (https://doi.org/10.6084/m9.figshare.28836158). This is in contrast with a score of 0 at both the N- and C-termini. In the second scoring function, for each position, we consider the scores of neighboring amino acids. To do so, we average the minimum scoring features using a sliding window of 7 residues, except at the three terminal residues where 4-6 residues were used (Fig 2C). This approach prevents us from selecting a tag site that is close to a less favorable region. The average minimum feature scores at S181 and S247 then are 0.19 and 0.18 respectively. By considering the raw per-residue scores and the average minimum score, we identified the S181 and S247 positions as most suited for tag insertion (Fig 2C,D).

We then sought to experimentally validate the efficacy of EpicTope’s predicted tagging sites for Smad5. To test both whether tagging Smad5 at the predicted sites preserves protein function and whether a tag is accessible to antibody binding, we created two constructs that inserted V5 tags into Smad5 at the top two predicted sites (S181 and S247). For comparison, we also created constructs with V5 tags inserted at the N- and C-terminal ends of the protein, regions that are better conserved and more ordered than S181 and S247 (Fig. 2). S181 and S247 are at the start and end of the linker region of Smad5, a disordered region (Fig. 2D), which has CDK8/9 phosphorylation sites for MAP kinases (Kretzschmar et al. 1999). The linker region connects the well-conserved MH1 and MH2 domains (Fig. 2C). The MH1 domain is essential for nuclear localization and DNA binding, while the MH2 domain mediates a slew of protein interactions such as receptor association and Smad-Smad binding (Derynck and Budi 2019, Macias et al 2015). Disrupting any of these core functions interferes with the ability of Smad5 to transduce BMP signaling (Macias et al 2015).

### EpicTope-predicted Smad5 tags preserve protein functionality

To test whether the S181-V5 and S247-V5 Smad5 proteins are functional, we performed a rescue experiment by injecting mRNA encoding these Smad5-V5 tags into zebrafish embryos deficient for Smad5. Smad5 transduces the BMP signaling protein, which functions in a concentration-dependent manner as a morphogen, patterning cells along the dorsal-ventral embryonic axis of all vertebrates during late blastula and gastrula stages of development (Greenfeld et al., 2021; Madamanchi et al., 2021; Zinski et al., 2017). Zebrafish dorsal-ventral axial patterning by BMP signaling has been extensively studied. Embryos with progressively reduced BMP signaling show progressively greater degrees of dorsalization due to the loss of ventral tissue fates and expansion of dorsal ones (Mullins et al., 1996; Mintzer et al., 2000; Tuazon et al., 2020). The early onset and progressive nature of this BMP-dependent process makes it ideal for performing quantitative rescue experiments. Here we used the *smad5^dtc24^* allele that carries a dominant-maternal antimorphic (dominant negative) mutation, which encodes an altered amino acid in the L3 loop of the Smad5 MH2 domain that mediates Smad-Smad interaction (Hild et al., 1999) (Fig 3A). The *smad5* transcript is maternally provided to eggs and hence heterozygous mothers carrying the *smad5^dtc24^*allele produce 100% strongly dorsalized embryos that are severely deficient in BMP signaling activity (Kramer et al., 2002; Mullins et al., 1996) (Fig. 3B). We refer to these as maternally mutant *smad5^dtc24^* embryos or M-*smad5^dtc24^* embryos. Though the *smad5^dtc24^* allele is antimorphic, it can be rescued by wild-type (WT) Smad1/5 (Hild et al., 1999; Nguyen et al., 1998).

**Figure 3:**
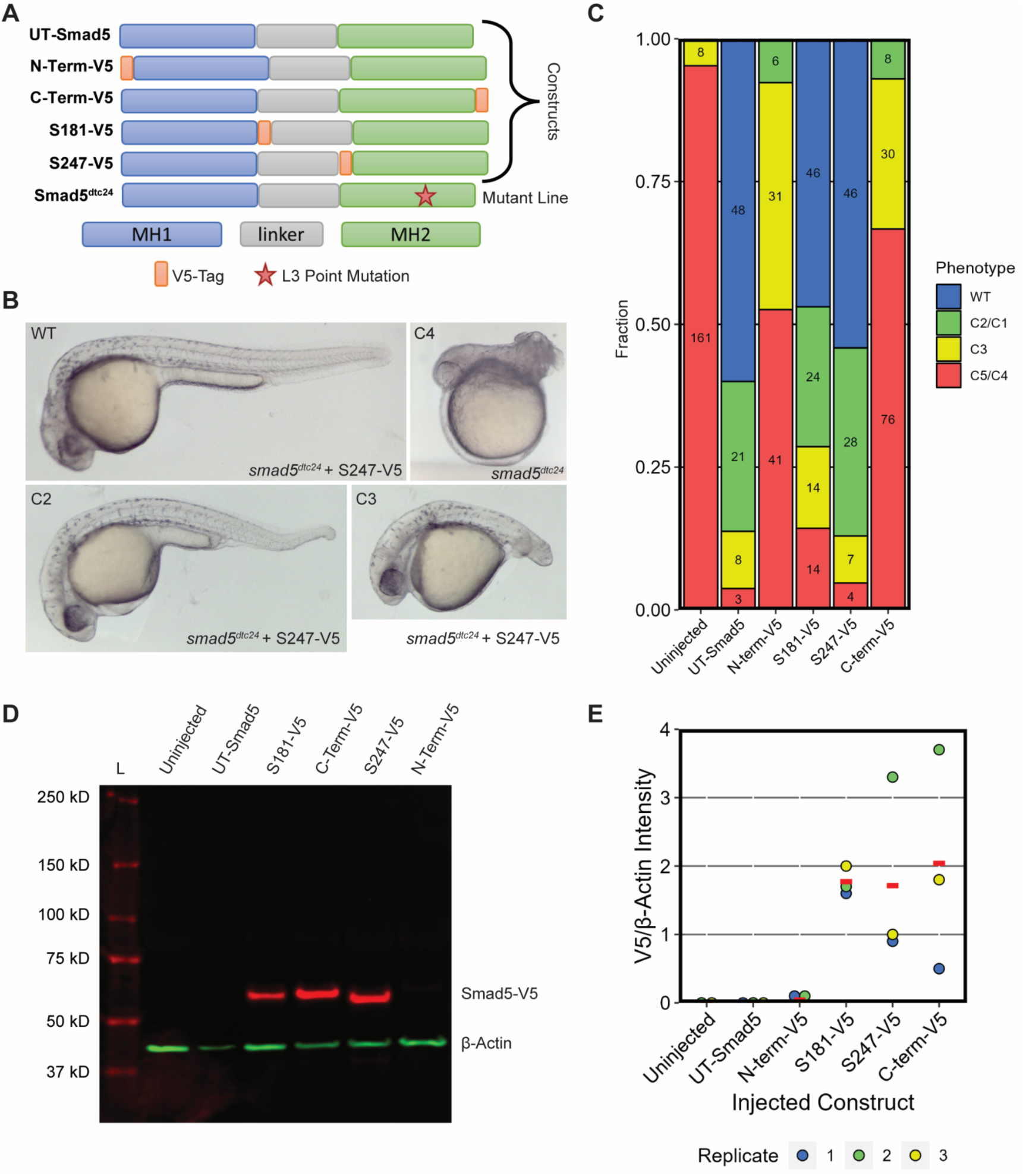
Internally V5 Tagged Smad5 Rescues Smad5 Mutant Embryos. A) Domain map of single V5 tagged constructs, untagged (UT) Smad5, and mutant Smad5. B) Representative images of 30 hpf M-*smad5^dtc24^* embryos injected with 150 pg of *smad5-S247-V5* RNA. C) Quantification of M-*smad5^dtc24^*embryos injected with 150 pg of UT or V5-tagged *smad5* RNA. The dorsalized classes C1-C5 shown in panel B are the standardized scoring scale used from Mullins et al, 1996. D) A western blot of extracts from M-*smad5^dtc24^* embryos injected with V5-tagged or untagged constructs and probed with anti-V5 and anti-β-Actin antibodies. Extracts of seven 6 hpf (shield stage) embryos were run in each lane. L is the molecular weight ladder. E) Quantification of western blots from 3 biological replicates of injections, Replicate 1 (RNA batch 1) and Replicates 2,3 (RNA batch 2) were from different RNA synthesis reactions. The red dash is the mean.

To test the functionality of our Smad5 constructs, we injected M-*smad5^dtc24^* embryos with mRNA encoding the single V5 tags at S181-V5, S247-V5, the N- or C-terminus, or untagged (UT)-*smad5*. We assessed the degree of phenotypic rescue at 30 hpf. We evaluated the level of dorsalization using the scoring scale from Mullins et al., 1996. Both the N- and C-terminally tagged Smad5 showed minimal rescue, consistent with these tag locations disrupting Smad5 functionality (Fig. 3B,C, S2). In contrast, both S181-V5 and S247-V5 Smad5 rescued embryos comparably to untagged Smad5 (Fig. 3B,C, S2). This shows that the internally-V5-tagged Smad5 proteins are similarly functional to WT Smad5 protein, while the N- and C-terminally tagged Smad5 are not.

To test whether the V5 tag interferes with protein stability, we performed western blot analysis on early gastrula embryos expressing, via mRNA injection, untagged or the V5-tagged Smad5 proteins. We found that the V5-tagged Smad5 protein was present at similar levels in the S181-V5, S247-V5, and C-terminal-V5 Smad5 expressing embryos, whereas only a faint band was evident in the N-terminal-V5 Smad5 expressing embryos (Fig. 3D, Fig. S3). When quantified relative to the β-Actin control, all V5-tagged Smad5 proteins were consistently present, while no V5-tagged protein was detected in the uninjected and UT Smad5 control conditions (Fig. 3E), as expected. This shows that S181-V5, S247-V5, and C-terminally V5-tagged Smad5 proteins are stable and present in injected embryos from *smad5^dct24^* females. Meanwhile N-terminally V5-tagged Smad5 is much less abundant, possibly due to misfolding and/or a stability issue caused by the N-terminal tag.

### Triple V5-tagged Smad5 is functional and effective for immunoprecipitation

To test if triple V5 (3xV5) tagging of Smad5 would disrupt protein functionality, we created additional constructs with 3xV5 tags at the previously described top predicted sites (Fig. 4A). We injected M-*smad5^dtc24^* embryos with S181-3xV5, S247-3xV5, or UT-*smad5* mRNAs and evaluated rescue of dorsalization at 30 hpf (Fig. 4B). Both the S181-3xV5 and S247-3xV5 proteins rescued embryos to ratios comparable to UT-*smad5* (Fig. 4C). Furthermore, the 3xV5 Smad5 tagged proteins were readily detectable in western blots (Fig. 4D). These results demonstrate that increasing from 1xV5 to 3xV5 internally-tagged Smad5 did not alter Smad5 functionality.

**Figure 4:**
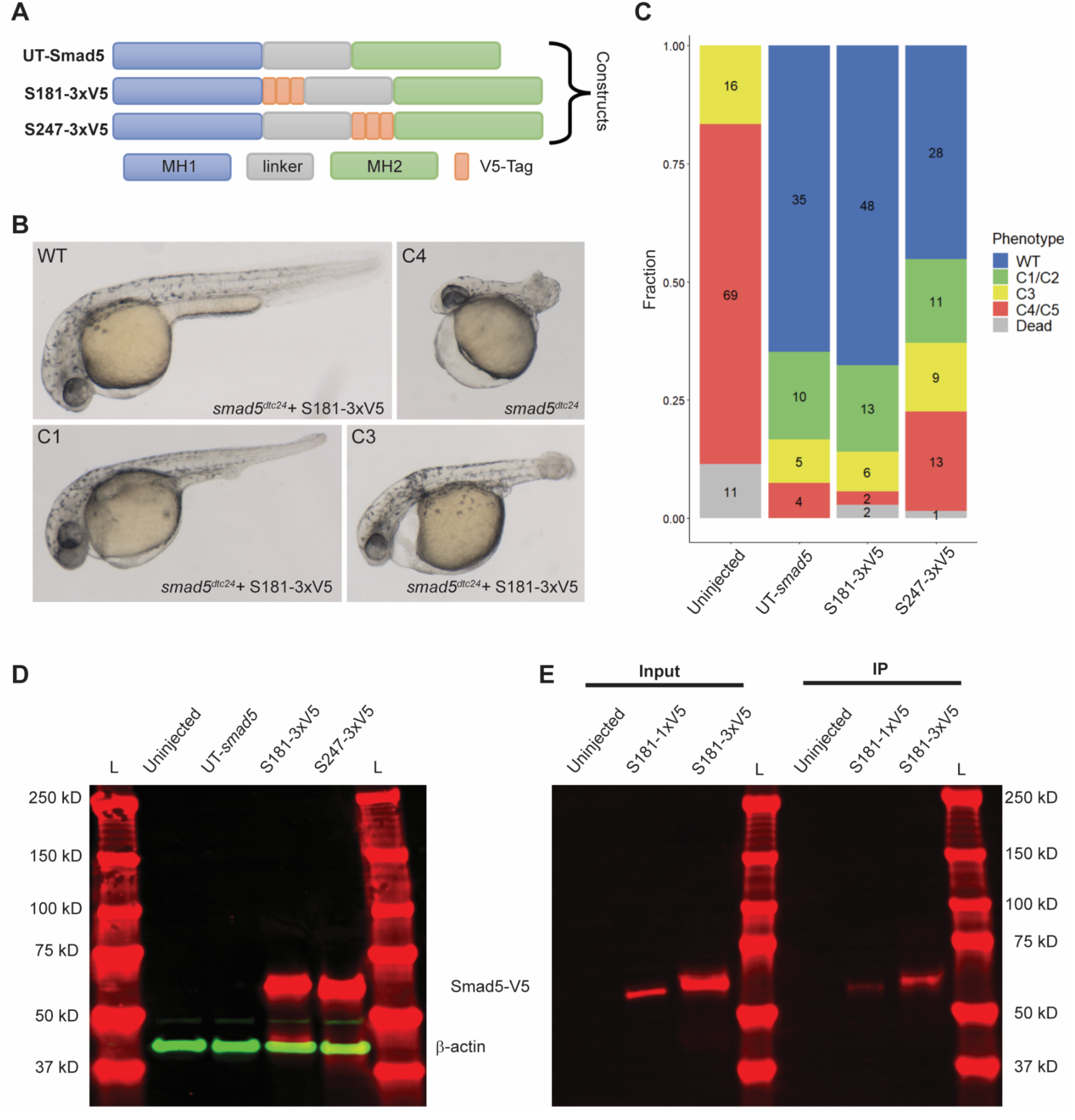
Triple V5-tagged Smad5 is functional in *smad5* mutant embryos and for immunoprecipitation. A) Domain map of UT-Smad5 protein and triple V5 (3xV5)-tagged Smad5 constructs. B) Representative images of 30 hpf M-*smad5^dtc24^* embryos uninjected or injected with 150 pg of *smad5-S181-3xV5*. C) Phenotypic quantification of M-*smad5^dtc24^* embryos injected with 150 pg of untagged (UT) or 3xV5-tagged *smad5* RNA from 2 biological replicates of injections. D) A western blot using anti-V5 (red) and anti-β-Actin (green) antibodies of M-*smad5^dtc24^* embryos injected with 3xV5-tagged or UT-*smad5* mRNA. The equivalent of seven 6 hpf (early gastrula) embryos were run in each lane. Representative of two biological replicates. E) A western blot of 6 hpf embryo extracts (input) and immunoprecipitated (IP) Smad5-S181-1x or −3xV5 protein. L is molecular weight ladder. Representative of two biological replicates.

Finally, we investigated whether the 3xV5 and 1xV5 tags in Smad5 could be used to immunoprecipitate tagged Smad5. We injected M-*smad5^dtc24^*embryos with S181-3xV5 or S181-1xV5 mRNA and immunoprecipitated V5-tagged protein from early gastrula (6 hpf) embryo extracts. Western blotting demonstrated that both the 1xV5 and 3xV5 tags at S181 can be readily immunoprecipitated with anti-V5 antibody. The 3xV5 tag yielded a stronger signal than the 1xV5 tag, likely due to the greater sensitivity conferred by the repeated tag. This shows that either single or triple V5 tags at the top two sites predicted by EpicTope in Smad5 are also suitable for immunoprecipitation.

### EpicTope-Predicted Smad5 Tags are Accessible in Embryos by Immunofluorescence

We then tested whether a single V5 tag inserted at S181 or S247 is sufficient to detect the subcellular localization of Smad5 by immunofluorescence. As in Fig 3, we injected M-*smad5^dtc24^* embryos with S181-V5, S247-V5 or UT-*smad5* mRNA. We immunostained embryos for the V5-tag, Phospho-Smad5 (P-Smad5), and Sytox Green (a DNA marker) and imaged them on a confocal (Zinski et al., 2017, 2019). Consistent with the rescue results shown in Figure 3C, we observed a WT-like gradient of P-Smad5 in UT, S181-V5, and S247-V5 Smad5 expressing embryos but not in uninjected, C- or N-terminally tagged Smad5 embryos (Fig 5A-F”). Consistent with the western blot analysis (Figure 3D), we observed V5-tagged protein in the S181, S247, and C-terminal V5-tagged Smad5 injected embryos but not in uninjected or N-terminal V5-tagged Smad5 injected embryos (Fig. 5A’”-F’”).

**Figure 5:**
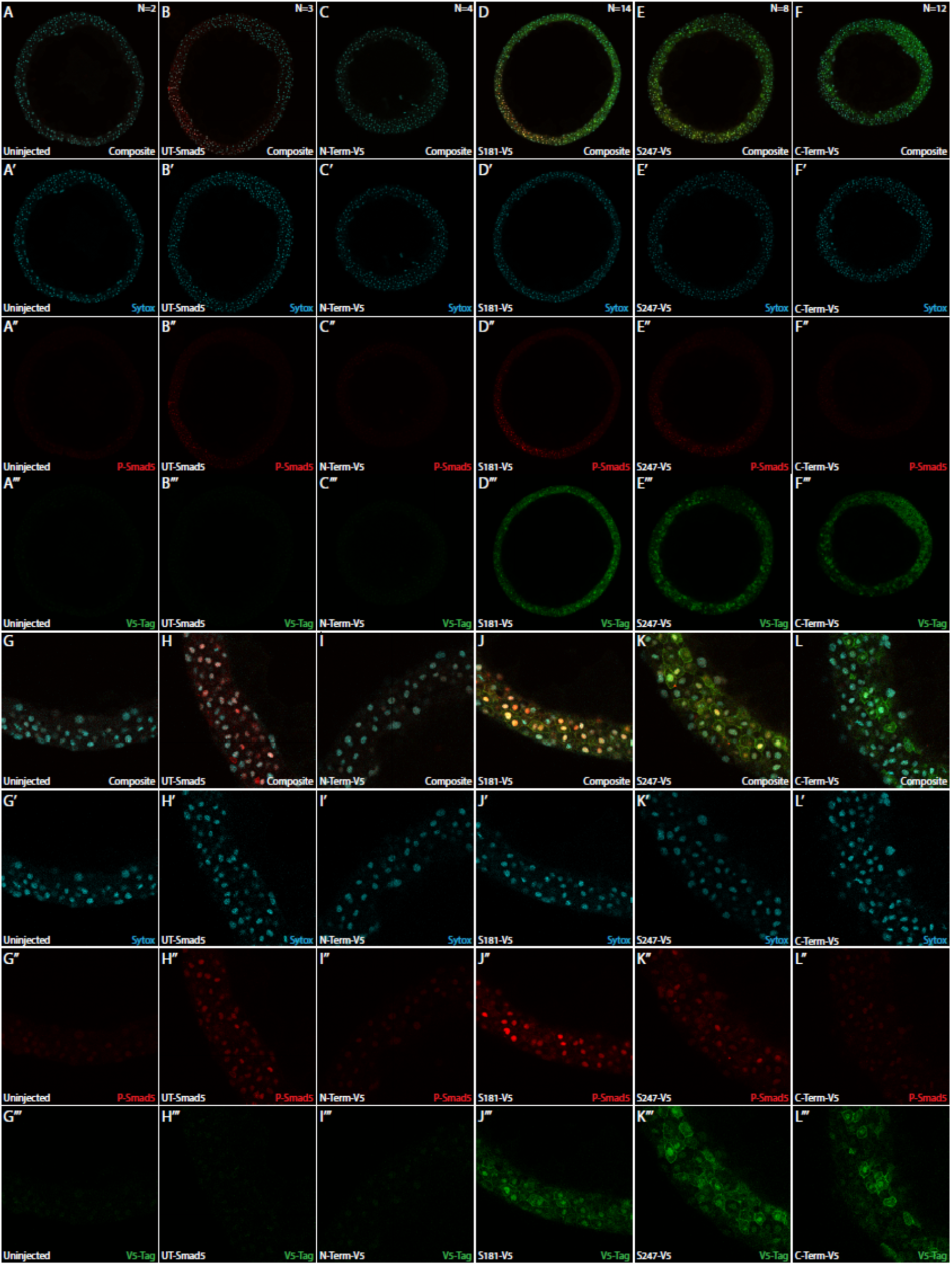
Internally tagged Smad5 colocalizes with nuclear phospho-Smad5. M-*smad5^dtc24^* embryos were injected with 150 pg of V5-tagged or untagged Smad5 and fixed at 5.7 hpf (early gastrula, germ ring stage). Embryos were immuno-stained for V5 (green), P-Smad5 (red), and for Sytox Green (DNA, blue). (A-F) Representative 25x zoom, 560 um X 560 um X 2.2 um confocal slices near the margin of the embryo. (G-L) Zoomed in 140 um X 140 um X 2.2 um sections of the embryos shown in A-F.

Receptor Smad proteins such as Smad5 reside in the cytoplasm until phosphorylated by a TGFβ Type I receptor, when they rapidly accumulate within the nucleus, activating BMP target gene expression (Hill, 2009; Schmierer & Hill, 2005). We sought to determine if V5-tagged Smad5 also localizes to the nucleus in the ventral region of injected embryos where extracellular BMP ligand concentrations should be high. Nuclear P-Smad5 was clearly evident in ventral nuclei of UT, S181-V5, and S247-V5 injected embryos, but only faintly present in nuclei of uninjected, N-term-V5, or C-term-V5 injected embryos (Fig. 5G-L”). V5-tagged Smad5 showed membrane, cytoplasmic, and nuclear localization in S181-V5, S247-V5, and C-term-V5 injected embryos (Fig. 5J’”-L’”). These results show that the internally-tagged V5 domain is effective for immunofluorescence microscopy.

### EpicTope prediction for Hdac1

We next tested EpicTope on a second protein, the chromatin regulator Histone deacetylase 1 (Hdac1), an essential, nuclear localized protein with a defined enzymatic domain that functions to deacetylate lysine residues in histone and non-histone targets. Hdac1 shows a high level of amino acid sequence conservation across species that decreases in the C-terminal region following the conserved histone deacetylase domain (Fig 6A). We used EpicTope to calculate Shannon entropy, secondary structure, RSA, and DBR feature scores for each amino acid of Hdac1. EpicTope predicted that the C-terminal region is relatively disordered (Fig 6B, teal line). Overall, the Hdac1 C-terminal disordered region scored well across all four parameters, having the highest RSA score, followed by internal amino acid A434 (Fig 6C). Alphafold2 predicts that the Hdac1 lysine deacetylase domain extends from amino acid 29 to 318 (Fig 6D), which places the A434 position within the C-terminal disordered region. The raw score is 0 and 0.24 for the N-terminus and A434, respectively, indicating that the N-terminus is a suboptimal location for tag insertion (https://doi.org/10.6084/m9.figshare.28836158).

**Figure 6.**
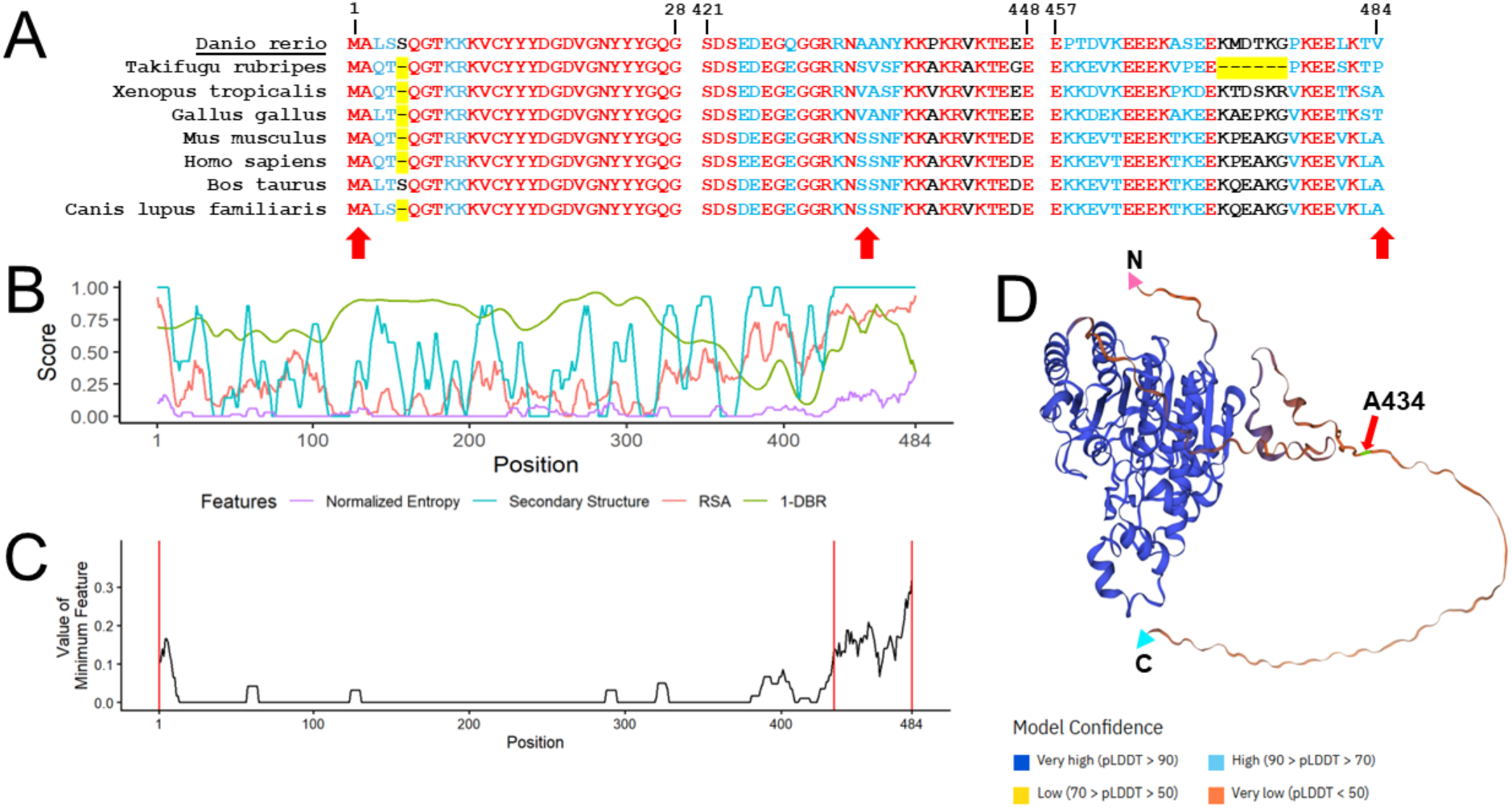
Hdac1 EpicTope features and prediction. EpicTope predictions for Hdac1, UniprotID: A0A2R8QIW0. A) MUSCLE multiple sequence alignment for Hdac1 positions 1-28, 421-448, and 457-484. Amino acids that are identical (red) or different (blue) between species. Red arrows indicate locations of V5 tag inserts, position indices are labeled in reference to zebrafish *Danio rerio*. B) Shannon entropy, secondary structure, relative solvent accessibility (RSA) and disordered binding region (DBR) features across the protein used in EpicTope prediction. Features are normalized to a 0-1 scale. C) Minimum feature values at each position are averaged over a sliding window of 7 residues, except for the three terminal residues where 4-6 residues are averaged and plotted. Red vertical lines indicate positions chosen for epitope tagging (N-term, A434, C-term). D) Hdac1 AlphaFold2 predicted structure; red arrow indicates internal tagging site A434, highlighted in green. C and N termini are labeled with blue and pink triangles, respectively.

### EpicTope-Predicted Hdac1 Tags Preserve Protein Functionality

To test whether Hdac1 proteins tagged at predicted optimal locations were functional, we performed transient rescue experiments of *hdac1* mutant zebrafish embryos. Zebrafish *hdac1* mutants display pleiotropic phenotypes in the developing nervous system that include microcephaly, microphthalmia, retinal coloboma, and defective axial extension (Henion et al., 2007; Ignatius et al., 2013; Schultz et al. 2018). For the functional assay we specifically measured the degree of rescue of the *hdac1* mutant axial extension phenotype (Fig. S4).

mRNAs were synthesized encoding positive control untagged *hdac1* (UT-Hdac1), *hdac1-A434-V5* (A434-V5), *hdac1-C-terminal-V5* (C-term-V5), and *hdac1-N-terminal-V5* (N-term-V5) (Fig. 7A). *hdac1* mutant embryos were generated by a cross between females heterozygous for the *hdac1^is70del4^*4-base pair frameshift allele (Schultz et al., 2018) and males carrying a 2A-mRFP loss of function knock-in allele that causes transcriptional termination and mRFP expression, *hdac1^is65off^* (Supplemental Fig. S4). One-cell stage embryos were injected with mRNAs and at 4 dpf all mRFP positive larvae were scored and placed into one of four categories indicating the degree of axial extension rescue (Fig. 7B). mRFP positive transheterozygous *hdac1 ^is70del4^l hdac1 ^is65off^* embryos were identified by PCR genotyping of the *hdac1^is70del4^* allele.

**Figure 7.**
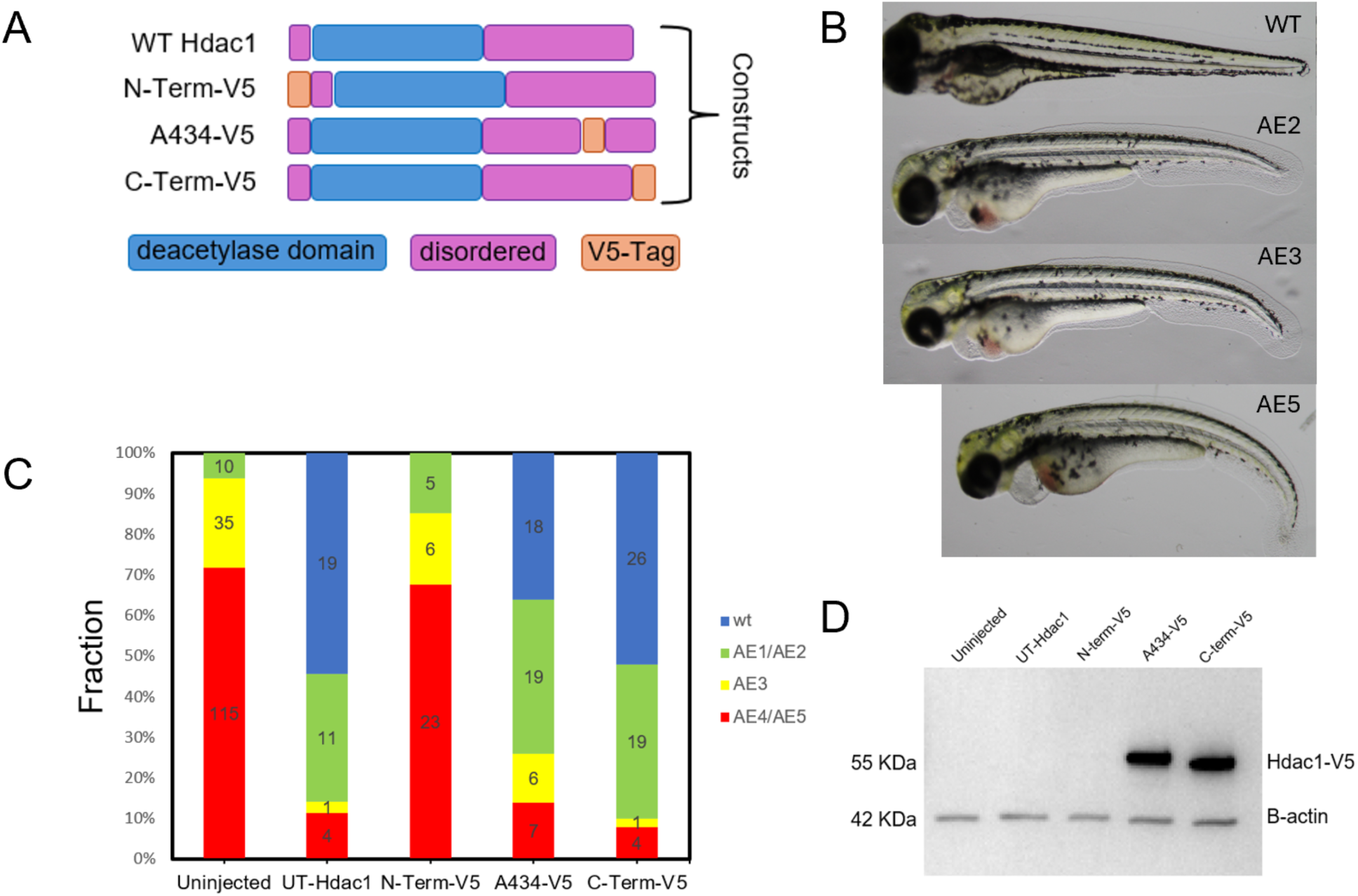
V5-tagged Hdac1 rescues *hdac1* null mutant embryos. A) Domain map of WT Hdac1, N-terminal, A434, and C-terminal V5 tag mRNAs. B) Scoring classification of embryos displaying a range of axial extension (AE) defects in transheterozygous *hdac1^is70del4^l hdac1 ^is65off^*4 dpf zebrafish embryos. C) Quantification of AE rescue of *hdac1^is70del4^l hdac1 ^is65off^* embryos injected with 50 pg of untagged or V5-tagged Hdac1 mRNA. Phenotypic classes were determined by trunk curvature and trunk-to-body ratio. D) Western blot probed with anti-V5 and anti-β-Actin antibodies of extracts from WT embryos injected with 50 pg of untagged or *V5*-tagged *hdac1* mRNA. Ten 6 hpf (shield stage) embryos were pooled for each lane.

*hdac1* C-term-V5 mRNA showed levels of rescue similar to WT UT-Hdac1 (Fig.7C). A434-V5 mRNA was able to rescue at comparable rates to C-term-V5 (Fig. 7C). However, the N-terminal V5 mRNA did not show any rescue (Fig.7C). These results are consistent with the EpicTope predictions and provide *in vivo* evidence that V5 integration at the Hdac1 C-terminus and internal A434 site are optimal locations for producing a functionally WT protein. These results are also consistent with previous studies using Hdac1 with a C-terminal FLAG tag for in vitro cell culture experiments (Pflum, 2001, Zhu, 2017). The absence of rescue by the N-term-V5 tag suggests that the insertion at the N-terminus abrogates Hdac1 function.

To determine the impact of the V5 tag on the stability of the tagged protein, we performed a western blot using extracts from 6 hpf WT embryos injected with each mRNA construct. Hdac1 protein tagged at A434 and at the C-terminus was stable and present at similar levels, whereas only a faint band was detected for the protein tagged at the N-terminus (Fig. 7D). Bands were not detected in extracts from un-injected embryos or embryos injected with untagged *hdac1* mRNA, demonstrating the specificity of the anti-V5 antibody (Fig.7D). This result indicates that the lack of rescue by the Hdac1 N-terminal V5 tag is due to either a defect in translation of the N-term-V5 mRNA or in the stability of the Hdac1-N-term-V5 tagged protein.

### EpicTope-predicted Hdac1 tags detected in embryos by immunofluorescence

We next tested whether rescue by the Hdac1-V5 tagged proteins correlated with nuclear localization visualized by whole-mount immunofluorescence in 6 hpf embryos. WT one-cell stage zebrafish embryos were injected with UT-*hdac1*, N-term-V5, A434-V5, or C-term-V5 *hdac1* mRNAs. Embryos were stained with anti-V5 and a nuclear stain (DAPI). High levels of V5 immunofluorescence were detected in embryos injected with the A434-V5 and C-term-V5 mRNAs (Fig. 8A). In contrast, N-term-V5 injected embryos showed very low levels of signal (Fig. 8A). Higher magnification revealed that both A434-V5 and C-term-V5 tagged Hdac1 were localized to the nucleus (Fig. 8B), consistent with the ability of these mRNAs to rescue the *hdac1* mutant phenotype. Although the level of N-terminal tagged protein was low, V5-Hdac1 protein also localized to the nucleus.

**Figure 8.**
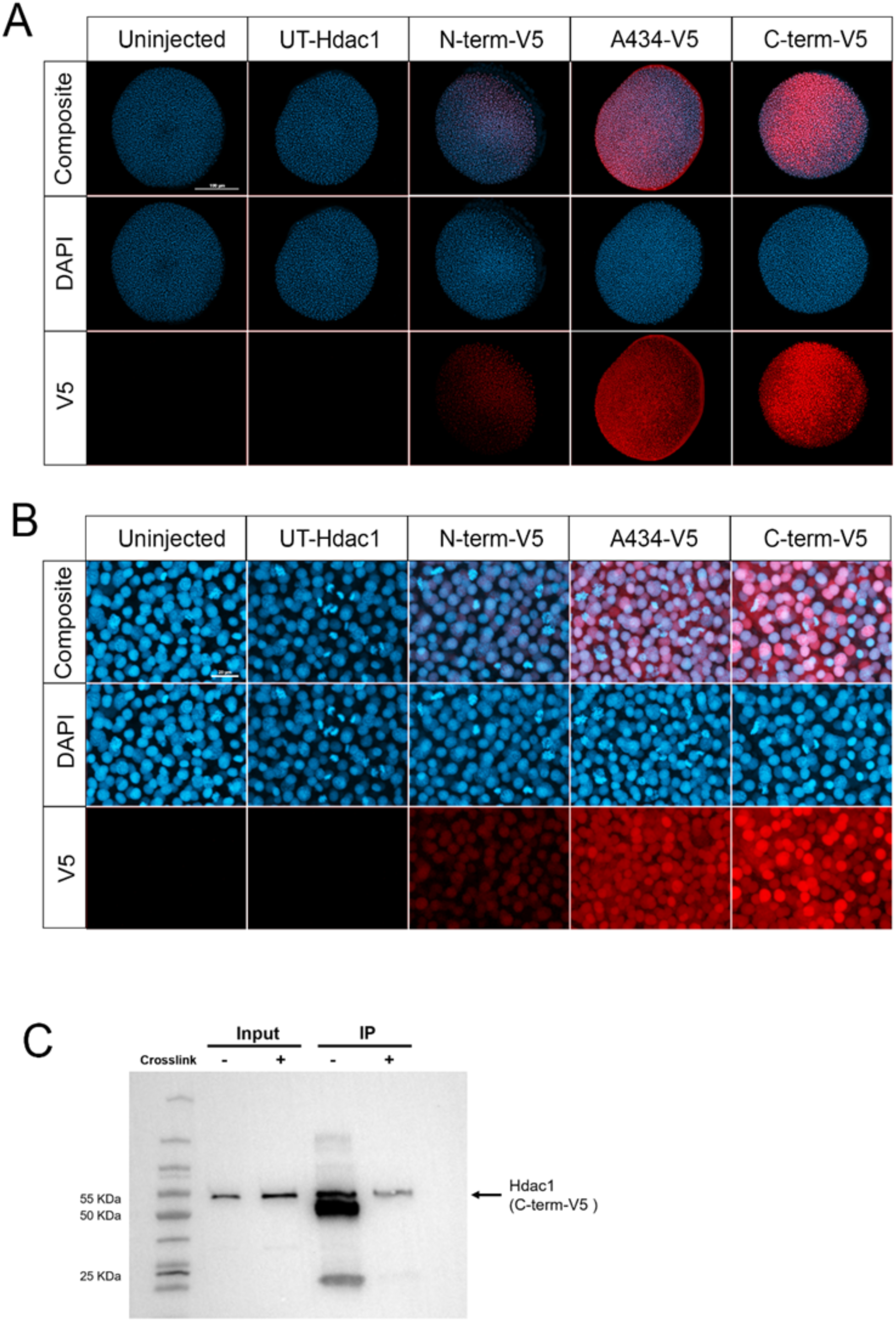
V5-tagged Hdac1 localizes to nuclei and can be immunoprecipitated. A) Confocal images of transheterozygous *hdac1^is70del4^l hdac1^is65off^*embryos injected with 50pg of UT or V5-tagged *hdac1* mRNA and fixed at 6 hpf (shield stage). Embryos were labeled with anti-V5 antibody (red) and stained with DAPI (blue). B) Representative higher magnification images showing nuclear localization. Scale bars: A 100 uM, B 20 uM. Experiments were performed in triplicate (n=9). C) Western blot of extracts (input) and immunoprecipitated (IP) C-term-V5 Hdac1 protein with (+) and without (-) antibody cross-linking to Protein G beads before IP. Without crosslinking the V5 antibody heavy (50 KDa) and light (25 KDa) chains are detected in the IP supernatant.

### A Single V5 tag is sufficient for Hdac1 protein capture

Lastly, we tested if a single V5 tag at the predicted optimal C-terminus location was sufficient for Hdac1-V5 immunoprecipitation. Extracts from WT embryos injected with 50 pg C-term-V5 mRNA were used for immunoprecipitation with anti-V5 antibody, with and without crosslinking to Protein G beads. Western blot of input extract and immunoprecipitated samples showed that the single C-terminal V5 tag precipitated Hdac1-V5 protein (Fig. 8C). Our results show that EpicTope can inform the design of multiple functional V5-tagged proteins suitable for immunolocalization and immunoprecipitation.

Together, our data show that EpicTope can predict epitope-tag insertion sites in a protein that do not interfere with protein function and are valuable tools to analyze protein localization by immuno-fluorescence microscopy, and to perform immunoprecipitation and western blots. We used the predictions to generate functional internally-tagged Smad5 and Hdac1 proteins. Introducing ALFA-tags to these predicted sites could enable applications ranging from single-molecule live imaging to cell labeling for FACS sorting (Boswell et al, 2025; Götzke et al., 2019; Girija et al., 2022; Westlund et al., 2023).

### Multiple scoring features impact insertion site predictions

In EpicTope analysis of the highly conserved Smad5 and Hdac1 proteins, entropy was a major predictor of tagging suitability (Figs. 2B,C, 6B,C). We applied the EpicTope workflow to five additional zebrafish proteins that are less highly conserved, namely, Bucky ball (Buc), Nanos1, Nanos3, Tdrd6, and Piwil1. Figure S5 shows that for these proteins, entropy is not the sole predictor of tagging suitability. The minimum feature value is driven by the secondary structure score and disordered region binding score for Buc in amino acid region 84-144 and 196-276, respectively (Fig. S5A). Nanos1 shows short dispersed amino acid regions (21-32, 47-67, 89-96) where secondary structure is the lowest of the 4 feature scores (Fig. S5B). Secondary structure and disordered region binding scores are the best predictor of tagging suitability in region 1-65 for Nanos3 (Fig. S5C). Similarly, secondary structure score guides tagging suitability for large regions of Piwil1 (Fig. S5D) and Tdrd6 (Fig. S5E). Thus, the multiple scoring features of EpicTope are indeed important to generate insertion site predictions for the large diversity of proteins encoded in the genome.

### EpicTope’s performance achieves equivalent precision as that of pathogenic variant predictors

We developed EpicTope to predict optimal tagging sites for proteins of interest. While our rescue experiments of zebrafish *smad5* and *hdac1* mutants showed that EpicTope-identified tag sites resulted in functionally viable proteins, we sought further to validate EpicTope on a larger set of possible targets. However, to the best of our knowledge, there is no large dataset of functionally negative and positive classifications of internally tagged proteins in zebrafish (or other organisms). We therefore considered the task of finding non-pathogenic, non-frameshifting insertion mutations to be a close analogous task. Following that rationale, we assessed EpicTope’s performance using a dataset of such human mutations collected from ClinVar (Landrum et al. 2017) and the Human Gene Mutation Database (Stenson et al. 2020). Both databases contain internal frameshift mutations associated with pathogenicity data. We compared EpicTope’s performance with that of two programs specifically designed for pathogenic protein prediction, Mutpred2 (Pejaver *et al* 2020) and SiftIndel (Hu and Ng 2013). See Methods, “Comparing EpicTope with MutPRED and SIFTindel”, for details.

We measured each method’s performance by calculating their F_1_ score, a standard score on a scale of 0-1 evaluating machine-learning models, as described in the Methods. SIFTindel achieved an F_1_ score of 0.694 (sd = 0.036), EpicTope an F_1_ score of 0.739 (sd = 0.035), and MutPred2 an F_1_ score of 0.831 (sd = 0.029) (Fig. S6). Thus, the predictive performance of EpicTope, when compared with SIFT-indel and Mutpred2, is favorable, with the EpicTope score lying between those of the two other predictors. We also note that pathogenicity only approximates protein function, as mutations can often cause disease by more complex mechanisms than simple loss-of-function (Taipale *et al*. 2019).

Following the overall comparison of the methods on a large dataset of known pathogenic insertion mutations, we compared EpicTope scores for *smad5* and *hdac1* with MutPred2, the latter having a higher average F_1_ score for predicting pathogenic non-frameshifting insertion mutants (Fig. S6). We used Mutpred2’s pathogenicity score as a proxy for identifying suitable epitope tag insertion sites, inverting the score to indicate the probability of non-pathogenic or non-disruptive insertion mutations. Mutpred2 predicted scores of 0.71 for the N-terminus and 0.63 and 0.64 for S181 and S247, respectively, thus not discriminating between functional internal tag positions and a non-functional end tag position. EpicTope scores align with our determined functional and non-functional Smad5 insertion sites (Fig. 3), assigning the N-terminus a score of 0, while 0.23 and 0.2 for S181 and S247, respectively (Fig. S7). For Hdac1, Mutpred2 assigned a score of 0.56 for the N-terminus and 0.64 for A434 (Fig. S8), which is similar to EpicTope’s relative scores for the two locations, 0 and 0.24, respectively (Fig. S8), consistent with our studies showing non-functional and functional sites, respectively, in Hdac1 (Fig. 7).

By design, EpicTope is more suited for identifying insertion tag sites than MutPred2 for two reasons. First, unlike MutPred2, EpicTope is designed to identify optimal tagging sites within a protein of interest and therefore assigns prediction scores to each position in the sequence.

Mutpred2 was not developed to make such predictions; rather, it uses mutant sequences as input, so to use MutPred2 to predict the functionality of insertion sites, like EpicTope, it requires generating a unique insertion mutant at each position of the protein of interest, a laborious task. The second fundamental difference is that EpicTope is designed to use on model organisms.

For example, the organisms over which sequence conservation is calculated may differ considerably depending on whether the protein of interest originates in mammals or, e.g., yeast. EpicTope allows the user to customize these features, adjusting the organisms for the sequence conservation calculation and the weights of individual features to best suit the user’s needs. Pathogenicity predictors like Mutpred2 and SiftIndel are trained on human pathogenic mutants and do not support any adjustment of the feature weights.

Our results show that EpicTope’s performance is comparable to that of MutPred2 and Siftindel, achieving an equivalent level of precision (Figure S4). Achieving a higher precision score is of a higher priority than recall, since the goal of tag insertion is to identify non-disruptive sites, and missing some correct sites can be tolerated. EpicTope achieved a slightly higher F_1_ score than SiftIndel but lagged behind Mutpred2. While Mutpred2’s F_1_ score was higher on average, we also show that Mutpred2 failed to discriminate between the known functional and non-functional tag insertion sites of Smad5 in our study. We believe this difference is due to the indirect relationship between protein function and pathogenicity. Pathogenicity is a complex emergent occurrence, often caused by defects in multiple genes or due to downstream pathways indirectly related to the function of a single gene. While Mutpred2 may be a better predictor of pathogenicity than EpicTope, MutPred2’s predictions do not necessarily translate to a more accurate prediction of protein function after tag insertion.

At the same time, the usage commonalities of these two programs justify their initial comparison: the underlying features or logic in Mutpred2 predictions could be integrated into EpicTope in the future. Mutpred2 predictions could also be directly included in a future version of EpicTope with proper weighting to tune the predictions more towards protein function.

### Limitations

While we have demonstrated that EpicTope correctly predicted four functional epitope-tag insertion sites in two proteins, and further predicted three less optimal sites in these same two proteins that we determined to be less or non-functional, future training of EpicTope combined with functional tests would further ensure its predictive power. Additionally, our functional tests were performed by expressing epitope-tagged proteins from injected mRNA into mutant embryos; genome editing of endogenous loci with these epitope tag insertions would further strengthen the predictive value of the EpicTope tool.

## Methods

### The EpicTope Tag Insertion Scoring Function

We used a scoring function based on four key protein features; sequence conservation, secondary structure, solvent accessibility, and disordered binding regions (Fig 1) to determine the ideal epitope tagging sites. A query protein is first identified by its UniprotID, and the sequence and AlphaFold2 predicted structure are retrieved through Uniprot’s API (Coudert et al., 2023). We determine the query’s sequence conservation by measuring the Shannon entropy (Eq. 1) at each position in a multiple sequence alignment (MSA) with homologous proteins (Shenkin et al., 1991).

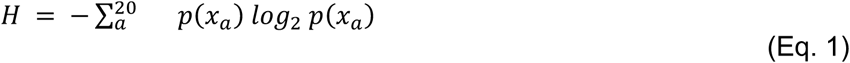

We identify homologs with BLAST, using the best hit match by lowest E value against a diverse set of vertebrate organisms; *Bos taurus, Canis lupus, Gallus gallus, Homo sapiens, Mus musculus, Takifugu rubripes, and Xenopus tropicalis* (Altschul et al., 1990), and then aligning these sequences with MUSCLE (Edgar, 2004). The Shannon Entropy (*H*) at each position is a function of the probability for a given amino acid (*a*) to appear at that position (*X) p*(*x_a_*) in the MSA, summed over all possible amino acids. We calculate secondary structures for the query protein using the Database of Secondary Structure of Proteins (DSSP), a tool for annotating secondary structure elements from protein structures (Kabsch & Sander, 1983). DSSP additionally provides an estimate of the solvent accessible surface area (*SASA*) for each position, and we calculate the relative solvent accessibility (*RSA*) by normalizing this estimate by the maximum solvent accessibility (SA_max_) for each amino acid (Eq. 2,(Tien et al., 2013)).

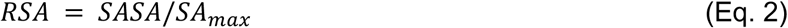

We retrieve estimates of the query protein’s predicted disordered protein-protein interaction regions from the IUPred2A web server, which uses ANCHOR2 for its prediction (Mészáros et al., 2018). After calculating and retrieving the key features, we normalize the values to a 0-1 scale. We then divide the Shannon entropy by 4.32, the maximum possible entropy, in bits, over 20 amino acids (log_2_(20) = 4.32). We bin the predicted secondary structure at each position (*X_s_*) into a numeric value (*SS*) based on expected sensitivity to tagging (Eq. 3).

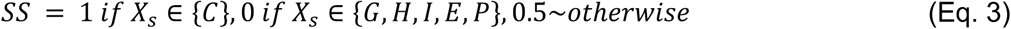

We expect defined structures such as alpha helices and beta strands (DSSP codes: *G,H,I, E, or P*) to be sensitive to disruption by inserted sequences, and assign these locations a score of 0. In contrast, when no secondary structure is assigned (*C*) it suggests that a tag insertion would be less disruptive to the structure. Therefore, C positions were assigned a score of 1. The ANCHOR2 disordered region binding score (*BR*) is the probability for a residue position to participate in a protein binding interaction. To avoid disrupting predicted protein interactions, we use 1 - BR as input in our scoring function.

We calculate a tagging score for each position *i* (*E_i_*), being the minimum value of all features (Eq. 4). We then seek out positions along the sequence where this minimum score is the highest. We use this approach to identify regions where all chosen features indicate tagging suitability. Calculating *E_i_* for every residue in a protein allows us to reject locations where insertions would be disruptive as indicated by at least one feature.

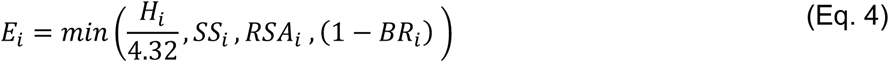

### Comparing EpicTope with Mutpred and SIFTindel

As a preliminary evaluation of EpicTope’s predictions, we compare the task of identifying a suitable epitope tag insertion site to predicting the pathogenicity of an in-frame insertion mutation. Although these two tasks are not fully equivalent, a mutated protein may still retain function even if it is disease-causing; assessing this task should provide a reasonable benchmark for EpicTope’s performance (Taipale et al. 2019).

We constructed a dataset of insertion mutations labeled pathogenic or benign. To do so we retrieved a dataset first compiled by Folkman et al. 2015, consisting of pathogenic and benign insertion-deletion mutants compiled from the Human Gene Mutation Database (Stenson et al. 2020). Data were retrieved in February 2024 from Varibench, a benchmark database comprising variation datasets for testing and training methods for variation effect prediction (Nair et al. 2012). We then filtered the data to include only insertion mutations of equivalent length to the V5 tag, resulting in 401 pathogenic or in-frame positive samples and 433 benign or negative samples. To update the Varibench dataset with new in-frame insertion mutants identified since their initial release, we also searched ClinVar, a database of human genetic variants with disease significance (Landrum et al. 2017). We cross-referenced the ClinVar results with VariBench and removed duplicate entries. Our final unique reference dataset consisted of 426 insertions labeled pathogenic and 446 labeled benign, summing to 872 samples total. (See https://doi.org/10.6084/m9.figshare.28836158 for complete dataset)

To comparatively assess EpicTope’s predictions, we fit a simple logistic regression model using EpicTope features to our reference dataset of pathogenic and benign in-frame insertion mutations. We modeled pathogenicity as a simple linear combination of EpicTope features; Shannon entropy, relative solvent accessibility, secondary structure score, ANCHOR2 disordered region binding score, the weighted sum of all scores (*S*_i_, Eq. 5), the minimum value score (*E_i_*), and each secondary structure individually represented as a one-hot encoding. We split the data into an 80:20 training and test set. We evaluated model performance by calculating the F1-score, or harmonic mean of the precision and recall.

The weighted sum of scores is calculated as the weighted sum of all features, with optional weights (*w*) for each variable, for each position *i* (Eq. 5).

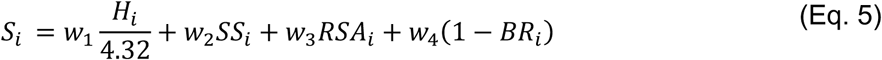

As an initial proof of concept, we set the Shannon entropy weight to 1.5 and all other weights to 1. These weights can be tuned to find an optimal contribution of each feature for ideal internal epitope tag insertion site prediction, and for now we used the weighted equation in the comparative assessment of EpicTope with MutPred and SIFTindel.

We then compared EpicTope’s performance with two similar predictors of indel impact, Mutpred2-indel and SIFT-indel. We note that SIFTindel does not give a probability score for pathogenicity, only returning pathogenic or benign labels. Therefore, precision-recall curve and receiver under operating characteristic (ROC) curve comparisons could only be made between Epictope and Mutpred2-indel. We also attempted a comparison with the DDIG-in model, but their web server was unavailable during testing (Folkman et al. 2015)

To directly assess Mutpred2’s pathogenicity predictions against EpicTope, we generated variants of Smad5 with the V5 inserted at every position in the sequence and predicted their pathogenicity with Mutpred2. We then compared the Mutpred2 prediction for each variant against the EpicTope score for the same position Mutpred2 scores predict pathogenicity, where a higher probability would indicate an insertion mutation more likely to disrupt function.

Therefore, we invert the predicted score by subtracting the score from 1, representing the probability the mutation is not disease-causing, to match EpicTope’s scale.

To compare the performance of the different methods we used common metrics for comparing prediction methods: precision, recall, and the F_1_ score defined as following:

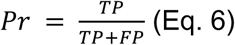

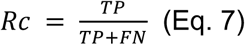

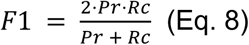

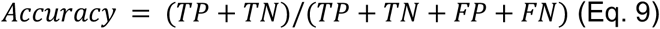

Where *TP*: the number of true positive predictions; *FP*: the number of false positives; *FN*: the number of false negatives. *Pr*: Precision; *Rc*: Recall; *F_1_*: the harmonic mean of precision and recall, which represents both precision and recall in one comparable metric. The predictions were benchmarked against known insertion mutations of comparable lengths to the tag we used, and that were documented experimentally as disease related.

### Zebrafish lines used in study

The WT TU line was used in all studies of Smad5, and the WT WIK strain, obtained from the Zebrafish International Resource Center (https://zebrafish.org/home/guide.php), was used in all Hdac1 studies. Fish were between 3 months and 1.5 years of age. Previously described zebrafish mutant lines used in this study: *smad5^dtc24^* (ID ZDB-ALT-980203-795, Mullins et al, 1996) and *hdac1^is70del4^* (Schultz et al., 2018). KBiosciences Competitive Allele-Specific PCR genotyping system (KASP) was used to genotype *smad5^dtc24^* with primers made by KBiosciences to the sequences: GTCAATTGATTTATTATTAGTAATATACACTTGTTGCTCTGAGTTTTAGATCAAGAACCAAGTGATATGAATATAATTCCATCCATCCATGTTTAATCTTCAACTTTTTTCTTTATTTCTTTCTATTCAAGGGTTGGGGTGCCGAGTACCACAGACAAGANGTGA[C/T]AAGCACCCCCTGCTGGATAGAAGTGCATCTCCACGGC CCCCTGCAATGGCTGGATAAAGTTCTAACACAAATGGGTTCCCCTCTGAACCCCATCTCTTCTGTCTCGTAATGATGGGCTGACCTGGGAGGAGCCT. Primers used for PCR genotyping of the *hdac1^is70del4^* allele: *hdac1* F 5’-GCAGACAGACATTGCCATTG-3’; *hdac1* R 5’-GCTCCAGAATGGCCAGTACA-3’, which specifically amplify the mutant allele and was visualized by gel electrophoresis. Isolation of the loss-of-function allele *hdac1^is65off^* is described below. Experimental protocols used in this study were approved by the University of Pennsylvania and Iowa State University Institutional Animal Care and Use Committee (IACUC-23-158, IBC-23-064) in compliance with American Veterinary Medical Association and the National Institutes of Health guidelines for the humane use of laboratory animals in research.

### *in vitro* gene expression construct assembly

The pCS2+ based vector for *in vitro* transcription of zSmad5, has been described previously (Kramer et al, 2002). Single-copy V5 epitope tags (GKPIPNPLLGLDST) were inserted by overlap extension PCR using Q5 High-Fidelity Polymerase (NEB #M0492S), amplifying the whole pCS2+ expression cassette from SP6 promoter (and primer binding site) to the T3 primer binding site downstream of the SV40 poly A site. PCR fragments were cloned and confirmed by Sanger sequencing of the whole amplicon. Triple V5 tags were introduced by adding to the 5’ of PCR primers as overlaps, PCR amplification of the whole plasmid and recircularization by Gibson Assembly (NEB #E2611S). Obtained clones were confirmed by Sanger sequencing of the insert between SP6 and T3 primers. Zebrafish *hdac1* cDNA was amplified from 500 ng total RNA isolated from 3 dpf embryos by RT-PCR using Superscript II (Invitrogen 11752) for first strand synthesis, followed by amplification with KOD polymerase (Sigma-Aldrich 71842). The 2008 bp *hdac1* cDNA was cloned into the pT3TS expression vector (Hyatt and Ekker, 1999) using HiFi cloning (Invitrogen A46624) of PCR products amplified with KOD (Sigma-Aldrich #71085). The V5 epitope tag was inserted into the *pT3TS-hdac1* vector using HiFi cloning (Invitrogen A46624) of PCR products amplified with KOD (Sigma-Aldrich #71085). Primers used for cDNA cloning and V5 epitope insertion are listed in Supplementary Table 1. Clones were confirmed by Sanger Sequencing (Fig. S9).

### Rescue with WT and V5-tagged mRNAs

Heterozygous *smad5^dtc24^* females were crossed to heterozygous males, which produce embryos with C4 or C5 dorsalized phenotypes (Mullins et al. 1996). Embryos from crosses between heterozygous *smad5^dtc24^* fish were injected at the 1-cell stage with 150 pg of mRNA. *smad5* mRNA was made from the N-terminal-V5, S181-V5, S181-3xV5, S247-V5, S247-3xV5, or C-terminal-V5 *smad5* plasmids generated for this paper or from a pCS2+ plasmid encoding untagged-Smad5, using the SP6 mMessage mMachine kit (Sigma Aldrich, AM1340). Rescue was assessed at 1 dpf using the scoring scale from Mullins et al. 1996. *hdac1* expression constructs *pT3TS-UT-hdac1* (untagged control), *pT3TS-N-term-V5, pT3TS-A434-V5*, and *pT3TS-C-term-V5* were used for *in vitro* transcription from 1 ug linear plasmid DNA to generate capped, polyadenylated mRNA using T3 mMessage mMachine High Yield Capped RNA transcription kit (ThermoFisher AM 1348). mRNA was purified using the ZYMO RNA Clean & Concentrator-5 kit (ZYMO R1015) and eluted in RNase-free water. One-cell stage embryos from a cross between heterozygous *hdac1^is70del4^*/+ and *hdac1^is65off^* /+ adults were injected with 50 pg *hdac1* untagged and V5-tagged mRNAs and mRFP positive larvae were scored for axial extension rescue at 4 dpf. All larvae were PCR genotyped for the presence of the *hdac1^is70del4^* 4 bp deletion allele to identify control *hdac1^i65off^/+* and transheterozygous *hdac1^is70del4^lhdac1^i65off^* individuals.

### Immunolabeling, microscopy, and image acquisition

Zebrafish embryos were fixed at 6 hours post fertilization (hpf) in 4% PFA in PBS. We then performed double immunolabeling for the V5-tag and P-Smad5 to quantify nuclear localized P-Smad5 and Smad5-V5 using mouse anti-V5 (ThermoFisher R960-25) and rabbit anti P-Smad5 antibody (Cell Signaling Technology 13820), as described (Zinski et al., 2019). Hdac1-V5 immunolabeling was performed with the mouse anti-V5 antibody. Alexa Fluor 594 (ThermoFisher A-11005) secondary antibody was used at 1:500. Embryos were counterstained with 5 ug/ml DAPI (ThermoFisher D1306) for 10’ at room temperature. Embryos were imaged for immunofluorescence on a Zeiss LSM 880 laser scanning confocal microscope with LD LCI Plan-Apochromat 25×/0.8 Immersion Corr DIC M27 multi-immersion and Plan-Apochromat 63x/1.40 Oil DIC M27 objectives, or on a Zeiss LSM 800 laser scanning confocal microscope with Plan-Apochromat 10x/0.45 M27 and Plan-Apochromat 20x/0.8 M27 objectives. For live imaging, larvae were anesthetized in 0.015% Tricaine MS-222 Ethyl 3-aminobenzoate methanesulfonate (Sigma E10521) and mounted in 1.2% low melt agarose (Promega V2111) in embryo media. Larvae were imaged on a Leica M165 FC stereomicroscope and images captured with a Canon Rebel T3 camera using EOS Utility software (Canon).

### Immunoprecipitation

Embryo lysate preparation was adapted from the protocol outlined in Haws et al., 2023. For V5-tagged Smad5 immunoprecipitation, embryos were injected with 150 pg of S181-3xV5 or S181-1xV5 *smad5* mRNA at the 1-cell stage and 50 embryos were collected at 6 hpf. For V5-tagged Hdac1 immunoprecipitation, embryos were injected with 50 pg of UT or V5-tagged *hdac1* mRNA and collected at 6 hpf. Embryos were dechorionated and deyolked manually at 6 hpf and stored in pools of ten at −80 °C. All steps were performed as described (Haws et al, 2023) until sample lysis. A modified lysis buffer (50 mM pH 8.0 Tris HCl, 150 mM NaCl, 5 mM EDTA, Roche cOmplete Mini Protease Inhibitor Cocktail (Millipore Sigma 11836153001), 1% v/v Triton X-100) was used to lyse dissociated cells before maceration with a micropestle. The lysate was incubated on ice for 10 minutes before being centrifuged at 20,000g for 1 minute to pellet debris. 75 µL of the clarified lysate supernatant was then transferred and immediately used or snap-frozen on dry ice and stored at −80°.

For immunoprecipitation, Dynabeads Protein G magnetic beads (ThermoFisher 10009D) were washed 2x with wash buffer (50 mM pH 8.0 Tris HCl, 150 mM NaCl, 1 mM EDTA, 1% v/v Triton X-100). For Smad5 immunoprecipitation, the beads for each IP were incubated with 1 µg mouse monoclonal anti-V5 antibody (ThermoFisher R960-25) in wash buffer with 100 ng/µL BSA for 40 minutes at RT. Beads for Hdac1 immunoprecipitation were incubated with 2 µg mouse monoclonal anti-V5 primary in wash buffer for 10 minutes at RT. After incubation, the beads were washed 3x with wash buffer. For crosslinking, the beads were washed 2x with 0.2 M triethanolamine buffer (pH 8.2). The beads were then resuspended in fresh 20 mM DMP (dimethyl pimelimidate) in 0.2 M triethanolamine buffer and incubated with gentle mixing for 30 minutes at RT. Crosslinking was stopped by discarding the supernatant and incubating beads in 50 mM Tris (pH 7.5) for 15 minutes at RT. Crosslinked beads were washed 3x with PBST. For Smad5 immunoprecipitation, antibody-loaded beads were incubated with 50 µL of sample lysate per IP in wash buffer overnight at 4°. For Hdac1 immunoprecipitation, beads were incubated with the sample lysate in wash buffer at 0.35 µg/µL for four hours at 4 ^O^C. All beads were then washed 3x with wash buffer at RT. The Dynabeads-Ab-Ag complex was resuspended in 1x Laemmli Sample Buffer (BioRAD 1610737) and 5% beta-mercaptoethanol before being denatured at 95°C for 5 minutes. The remaining 25 µL of input sample lysate was also denatured with 1x Laemmli Sample Buffer. Western blots were performed as described below.

### Western Blot

Embryos were dechorionated and the yolks manually removed at 6 hpf. Samples were flash-frozen in liquid nitrogen and stored overnight at −80°C. Seven or ten embryos were pooled for each condition. Samples were denatured at 95°C for 5 minutes in 2X Laemlli buffer (65.8 mM Tris-HCl, pH 6.8, 2.1% SDS, 26.3% (w/v) glycerol, 0.01% bromophenol blue) and 5% beta-mercaptoethanol. Denatured protein in Laemlli buffer was loaded into a 4-15% gradient SDS-Page gel in a Mini Trans-blot cell (BioRAD) with running buffer (in dH2O, 25 mM Tris, 192 mM glycine, 0.1% SDS) and run at 50V for 10 minutes, followed by 100V for 80 minutes. Proteins were transferred to LF-PVDF membrane in transfer buffer (in dH2O, 20% methanol, 25 mM Tris, 192 mM glycine) at 4°C at 100V for 50 minutes. The membrane was washed 3x for 15 minutes in 1xTBST (150 mM NaCl, 10 mM Tris pH 8.0, 0.1% Tween20) or 1x PBST (137 mM NaCl, 10.1 mM Na2HPO4, 2.7 mM KCl, 2 mM KH2PO pH 7.2, 0.05% Tween20) then for 2 hours in 4% skim milk in TBST or PBST. The membranes were incubated with primary antibodies (1:1000 rabbit anti-β-actin (A2066 Sigma Aldrich), 1:2000 mouse anti-V5, or 1:2000 rabbit polyclonal anti-b-actin (Cell Signaling 4967s), 1:5000 mouse monoclonal anti-V5 (ThermoFisher R960-25)) in 4% milk in TBST or PBST O/N at 4°C. Membranes were then washed 3x for 15 minutes before being incubated in secondary (1:10000 anti-mouse DyLight 680 and 1:10000 anti-rabbit DyLight 800 or 1:5000 anti-mouse HRP Invitrogen 31430 and 1:5000 anti-rabbit HRP Invitrogen 31480) for 40 minutes at RT in the dark. Membranes were washed 3x in TBST or PBST and imaged on a LI-COR imaging system (LI-COR Biosciences) or Invitrogen iBright 1500 (ThermoFisher) imaging system.

All plasmids and fish generated or used are available upon request or can be found at the Zebrafish International Resource Center or Addgene.

## Data and resource availability

All relevant data and details of resources can be found within the article and its supplementary information.

## Acknowledgements

We thank the National Institutes of Health (NIH) and others for grant funding that supported these studies: NIH R24 OD020166 and NIH R24 OD036201 to I.F. and M.M., NIH R01GM145937 and SEED grant funding from the Translational AI Center at Iowa State University to I.F., NIH R35-GM131908 and 3R35GM131908-05S1 to M.C.M, and NIH R21 HD103982 to D.B.. We thank the University of Pennsylvania Cell & Developmental Biology Microscopy Core, especially Andrea Stout and AJ Lucy, for helpful support.

## Supplementary Materials

The raw data scores for Smad5 and Hdac1 proteins are provided on Figshare https://doi.org/10.6084/m9.figshare.28836158

The EpicTope software is available on GitHub, https://github.com/FriedbergLab/Epictope

The EpicTope source code can be found at Zenodo, https://doi.org/10.5281/zenodo.17957808

## References

Altschul, S. F., Gish, W., Miller, W., Myers, E. W., & Lipman, D. J. (1990). Basic local alignment search tool. Journal of Molecular Biology, 215(3), 403–410.

Boswell CW, Hoppe C, Sherrard A, Miao L, Kojima ML, Martino P, Zhao N, Stasevich TJ, Nicoli S, Giraldez AJ. (2025). Genetically encoded affinity reagents are a toolkit for visualizing and manipulating endogenous protein function in vivo. Nature Communications, 16(1):5503. doi: 10.1038/s41467-025-61003-w.

Burg, L., Zhang, K., Bonawitz, T., Grajevskaja, V., Bellipanni, G., Waring, R., & Balciunas, D. (2016). Internal epitope tagging informed by relative lack of sequence conservation. Scientific Reports, 6, 36986.

Coudert, E., Gehant, S., de Castro, E., Pozzato, M., Baratin, D., Neto, T., Sigrist, C. J. A., Redaschi, N., Bridge, A., & UniProt Consortium. (2023). Annotation of biologically relevant ligands in UniProtKB using ChEBI. Bioinformatics, 39(1). 10.1093/bioinformatics/btac793

Derynck, R., & Budi, E. H. (2019). Specificity, versatility, and control of TGF-β family signaling. Science signaling, 12(570). 10.1126/scisignal.aav5183

Edgar, R. C. (2004). MUSCLE: a multiple sequence alignment method with reduced time and space complexity. BMC Bioinformatics, 5, 113.

Folkman, L., Yang, Y., Li, Z., Stantic, B., Sattar, A., Mort, M., Cooper, D. N., Liu, Y., & Zhou, Y. (2015). DDIG-in: detecting disease-causing genetic variations due to frameshifting indels and nonsense mutations employing sequence and structural properties at nucleotide and protein levels. Bioinformatics, 31(10), 1599–1606.

Gibb N, Lazic S, Yuan X, Deshwar AR, Leslie M, Wilson MD, Scott IC. (2018). Hey2 regulates the size of the cardiac progenitor pool during vertebrate heart development. Development. 145(22). 10.1242/dev.167510

Götzke, H., Kilisch, M., Martínez-Carranza, M., Sograte-Idrissi, S., Rajavel, A., Schlichthaerle, T., Engels, N., Jungmann, R., Stenmark, P., Opazo, F., & Frey, S. (2019). The ALFA-tag is a highly versatile tool for nanobody-based bioscience applications. Nature Communications, 10(1), 4403.

Greenfeld, H., Lin, J., Mullins, M.C. (2021) The BMP signaling gradient is interpreted through concentration thresholds in dorsal-ventral axial patterning. PLoS Biology. 19:e3001059.

Hashiguchi, M., & Mullins, M. C. (2013). Anteroposterior and dorsoventral patterning are coordinated by an identical patterning clock. Development, 140(9), 1970–1980.

Hata, A., & Chen, Y.-G. (2016). TGF-β Signaling from Receptors to Smads. Cold Spring Harbor Perspectives in Biology, 8(9). 10.1101/cshperspect.a022061

Haws, W., England, S., Grieb, G., Susana, G., Hernandez, S., Mirer, H., & Lewis, K. (2023). Analyses of binding partners and functional domains for the developmentally essential protein Hmx3a/HMX3. Scientific Reports, 13, 1151.

Hild, M., Dick, A., Rauch, G. J., Meier, A., Bouwmeester, T., Haffter, P., & Hammerschmidt, M. (1999). The smad5 mutation somitabun blocks Bmp2b signaling during early dorsoventral patterning of the zebrafish embryo. Development, 126(10), 2149–2159.

Hill, C. S. (2009). Nucleocytoplasmic shuttling of Smad proteins. Cell Research, 19(1), 36–46.

Hu, Jing, and Pauline C. Ng. “SIFT Indel: predictions for the functional effects of amino acid insertions/deletions in proteins.” PloS one 8.10 (2013): e77940.

Hyatt, T. M., & Ekker, S. C. (1999). Vectors and techniques for ectopic gene expression in zebrafish. Methods in cell biology. Methods in cell biology, 59, 117–126.

Igreja, C., Loschko, T., Schäfer, A., Sharma, R., Quiobe, S. P., Aloshy, E., Witte, H., & Sommer, R. J. (2022). Application of ALFA-Tagging in the Nematode Model Organisms Caenorhabditis elegans and Pristionchus pacificus. Cells, 11(23). 10.3390/cells11233875

Kabsch, W., & Sander, C. (1983). Dictionary of protein secondary structure: pattern recognition of hydrogen-bonded and geometrical features. Biopolymers, 22(12), 2577–2637.

Kramer, C., Mayr, T., Nowak, M., Schumacher, J., Runke, G., Bauer, H., Wagner, D. S., Schmid, B., Imai, Y., Talbot, W. S., Mullins, M. C., & Hammerschmidt, M. (2002). Maternally supplied Smad5 is required for ventral specification in zebrafish embryos prior to zygotic Bmp signaling. Developmental Biology, 250(2), 263–279.

Kretzschmar, Marcus, et al. “A mechanism of repression of TGFβ/Smad signaling by oncogenic Ras.” Genes & development 13.7 (1999): 804–816.

Landrum, Melissa J., et al. “ClinVar: improvements to accessing data.” Nucleic acids research 48.D1 (2020): D835–D844.

Macias, M. J., Martin-Malpartida, P., & Massagué, J. (2015). Structural determinants of Smad function in TGF-β signaling. Trends in biochemical sciences, 40(6), 296–308. 10.1016/j.tibs.2015.03.012

Madamanchi, A., Mullins, M. C., & Umulis, D. M. (2021). Diversity and robustness of bone morphogenetic protein pattern formation. Development, 148(7). 10.1242/dev.192344

Mészáros, B., Erdos, G., & Dosztányi, Z. (2018). IUPred2A: context-dependent prediction of protein disorder as a function of redox state and protein binding. Nucleic Acids Research, 46(W1), W329–W337.

Mintzer, K.A., Lee, M.A., Runke, G., Trout, J., Whitman, M., and Mullins, M.C. (2001) lost-a-fin encodes a type I BMP receptor, Alk8, acting maternally and zygotically in dorsoventral pattern formation. Development (Cambridge, England). 128(6):859–869.

Mullins, M. C., Hammerschmidt, M., Kane, D. A., Odenthal, J., Brand, M., van Eeden, F. J., Furutani-Seiki, M., Granato, M., Haffter, P., Heisenberg, C. P., Jiang, Y. J., Kelsh, R. N., & Nüsslein-Volhard, C. (1996). Genes establishing dorsoventral pattern formation in the zebrafish embryo: the ventral specifying genes. Development, 123, 81–93.

Nair, Preethy Sasidharan, and Mauno Vihinen. “Vari Bench: A benchmark database for variations.” Human mutation 34.1 (2013): 42–49.

Nguyen, V. H., Schmid, B., Trout, J., Connors, S. A., Ekker, M., & Mullins, M. C. (1998). Ventral and lateral regions of the zebrafish gastrula, including the neural crest progenitors, are established by a bmp2b/swirl pathway of genes. Developmental Biology, 199(1), 93–110.

Oesterle, S., Roberts, T. M., Widmer, L. A., Mustafa, H., Panke, S., & Billerbeck, S. (2017). Sequence-based prediction of permissive stretches for internal protein tagging and knockdown. BMC Biology, 15(1), 100.

Pagel, K. A., Antaki, D., Lian, A., Mort, M., Cooper, D. N., Sebat, J., Iakoucheva, L. M., Mooney, S. D., & Radivojac, P. (2019). Pathogenicity and functional impact of non-frameshifting insertion/deletion variation in the human genome. PLoS Computational Biology, 15(6), e1007112.

Pejaver, Vikas, et al. “Inferring the molecular and phenotypic impact of amino acid variants with MutPred2.” Nature Communications 11.1 (2020): 5918.

Schmierer, B., & Hill, C. S. (2005). Kinetic analysis of Smad nucleocytoplasmic shuttling reveals a mechanism for transforming growth factor beta-dependent nuclear accumulation of Smads. Molecular and Cellular Biology, 25(22), 9845–9858.

Stenson, Peter D., et al. “The Human Gene Mutation Database (HGMD®): optimizing its use in a clinical diagnostic or research setting.” Human Genetics 139 (2020): 1197–1207.

Shenkin, P. S., Erman, B., & Mastrandrea, L. D. (1991). Information-theoretical entropy as a measure of sequence variability. Proteins, 11(4), 297–313.

Taipale, Mikko. “Disruption of protein function by pathogenic mutations: common and uncommon mechanisms.” Biochemistry and Cell Biology 97.1 (2019): 46–57.

Tien, M. Z., Meyer, A. G., Sydykova, D. K., Spielman, S. J., & Wilke, C. O. (2013). Maximum allowed solvent accessibilites of residues in proteins. PloS One, 8(11), e80635.

Tuazon, F.B., Wang, X., Andrade, J.L., Umulis, D., Mullins, M.C. (2020) Proteolytic Restriction of Chordin Range Underlies BMP Gradient Formation. Cell Reports. 32:108039.

Tucker, J. A., Mintzer, K. A., & Mullins, M. C. (2008). The BMP signaling gradient patterns dorsoventral tissues in a temporally progressive manner along the anteroposterior axis. Developmental Cell, 14(1), 108–119.

Westlund, E., Bergenstråle, A., Pokhrel, A., Chan, H., Skoglund, U., Daley, D. O., & Söderström, B. (2023). Application of nanotags and nanobodies for live cell single-molecule imaging of the Z-ring in Escherichia coli. Current Genetics, 69(2-3), 153–163.

Zinski, J., Bu, Y., Wang, X., Dou, W., Umulis, D., & Mullins, M. C. (2017). Systems biology derived source-sink mechanism of BMP gradient formation. eLife, 6. 10.7554/eLife.22199

Zinski, J., Tuazon, F., Huang, Y., Mullins, M., & Umulis, D. (2019). Imaging and Quantification of P-Smad1/5 in Zebrafish Blastula and Gastrula Embryos. Methods in Molecular Biology, 1891, 135–154.

Zordan, R. E., Beliveau, B. J., Trow, J. A., Craig, N. L., & Cormack, B. P. (2015). Avoiding the ends: internal epitope tagging of proteins using transposon Tn7. Genetics, 200(1), 47–58.

