## Supplemental Figures and Tables for "EpicTope: predicting and validating non-disruptive epitope tagging sites"

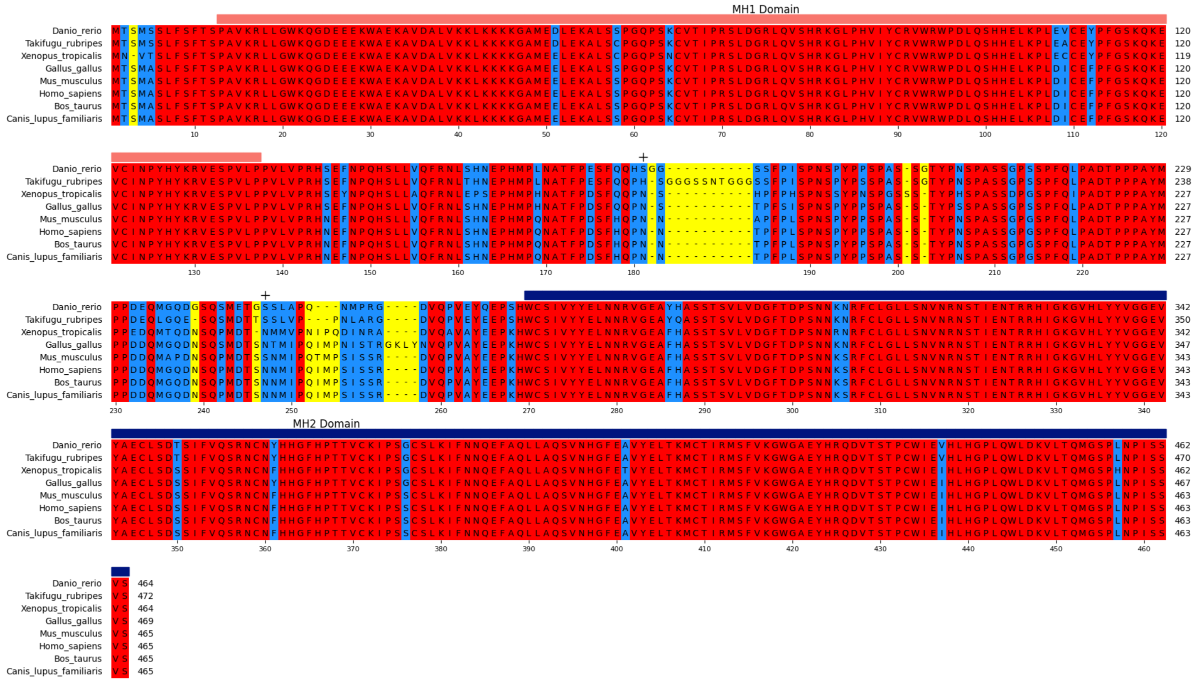

**Figure S1. Multiple Sequence Alignment for *Danio rerio* Smad5.** Amino acids identical between all species are highlighted in red and differences are highlighted blue. Amino acid length variation is shown in yellow. Location of internally inserted *Danio rerio* Smad5 tags at S181 and 247 are labeled with a black "+". MH1 and MH2 domains are indicated by horizontal red and dark blue bars, respectively. Total length of each Smad5 protein is shown at the end of the alignment, and position indices are labeled in reference to *Danio rerio*.

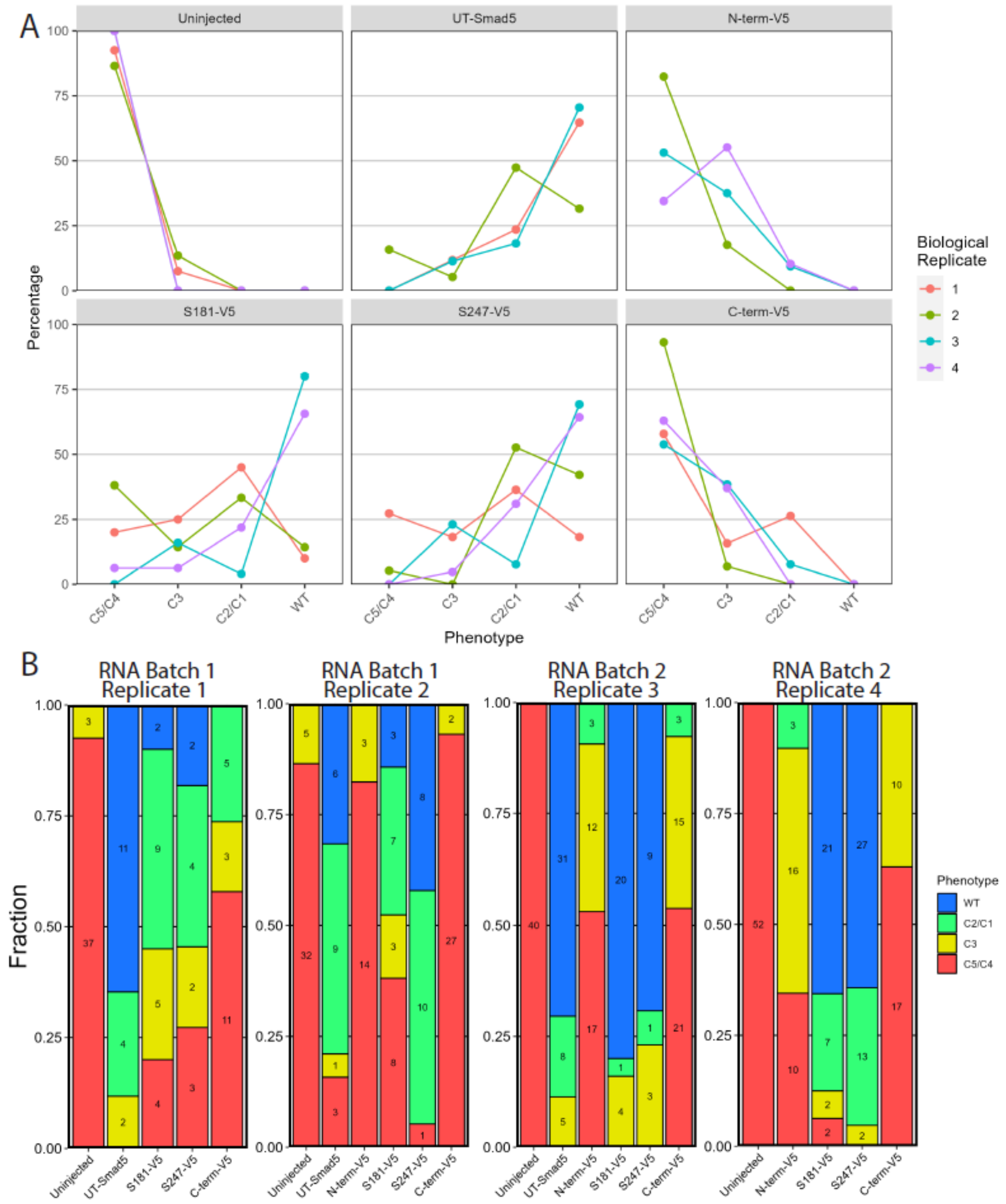


**Figure S2. Biological Replicates of V5-Tagged Smad5 Rescue.** Quantification of *smad5*$^{dtc24}$ mutant embryos injected with 150 pg of untagged (UT) or V5-tagged *smad5* RNA. The dorsalized classes C1-C5 are the scoring scale in Mullins et al, 1996. A) Points on the lines are the percent embryos with the phenotype specified on the x-axis. Different colored lines are different biological replicates. B) Bar graphs of data in A with the number of embryos noted in each bar.

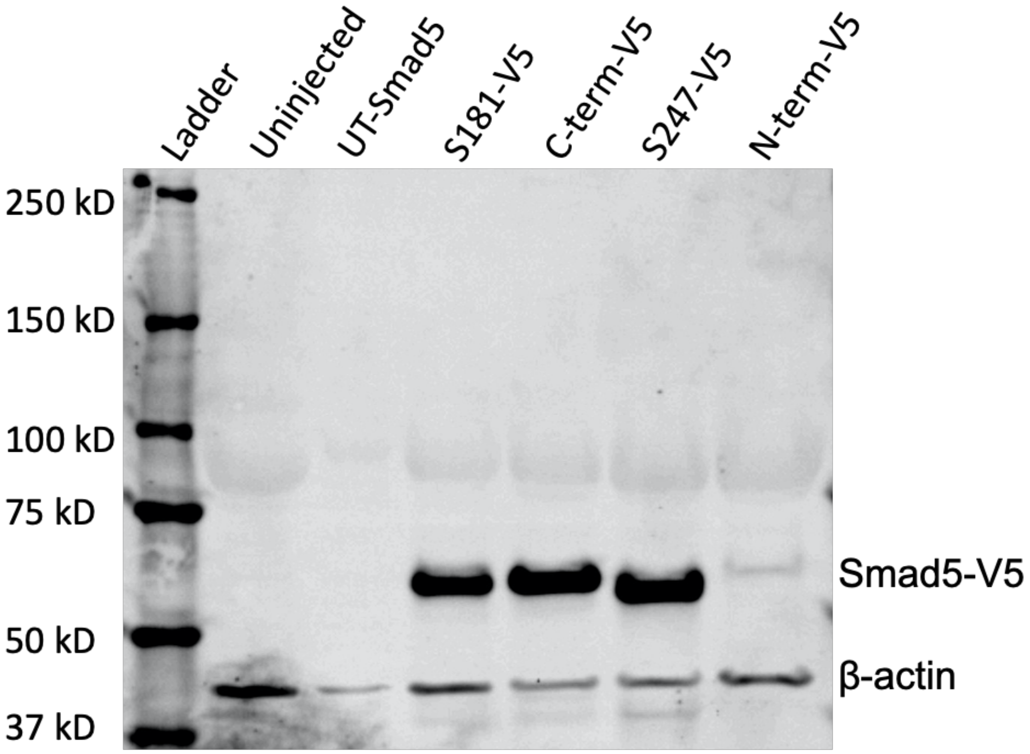


**Figure S3.** Longer exposure of the western blot in Figure 3 to show the N-terminal V5-tagged Smad5 band using anti-V5 and anti-β-Actin antibodies on extracts from embryos injected with V5-tagged or untagged (UT) mRNAs. Seven 6 hpf (shield stage) embryos were used in each lane.

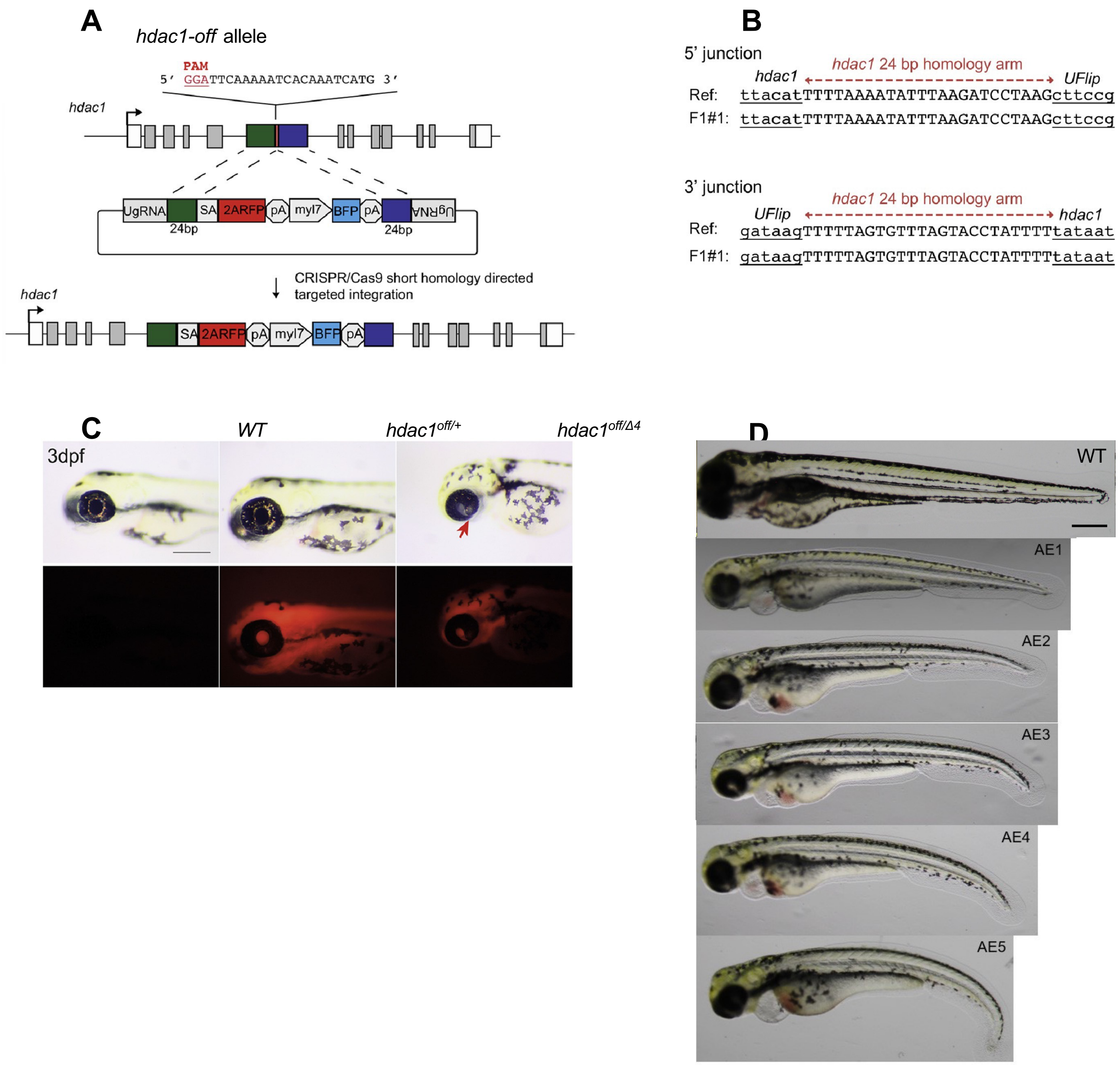


**Figure S4. Zebrafish loss of function *hdac1*$^{is65off}$ knock-in allele** A) Diagram of *hdac1*$^{is65off}$ loss of function KI allele. B) 5’ and 3’ genomic-UFlip cassette integration junctions were PCR amplified from F1 transgenic zebrafish fin clip genomic DNA. The PCR products were sequenced and aligned to the reference sequence expected for a precise integration at the genomic target site. C) *hdac1*$^{is65off}$ KI allele showing mRFP expression. Gross phenotype of 3 dpf transheterozygous *hdac1*$^{is70del4}$/*hdac1*$^{is65off}$ larva showing microcephaly and retinal coloboma (arrow). D) Embryos displaying all scoring categories show the range of axial extension defect phenotypes in transheterozygous *hdac1*$^{is70del4}$/*hdac1*$^{is65off}$ 4 dpf zebrafish embryos. Scale bars C,D 200 μM.

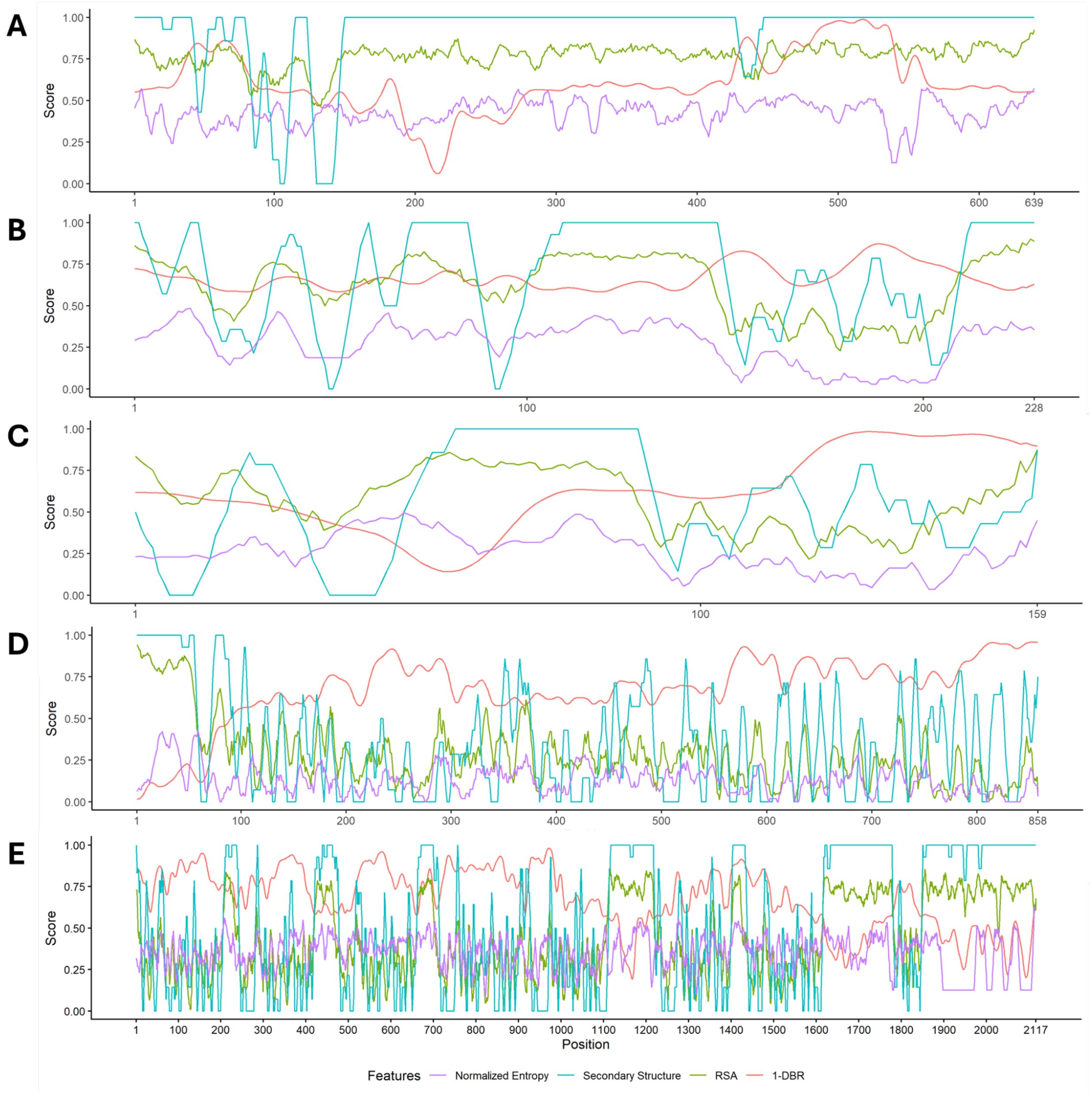


**Figure S5. EpicTope feature scores for additional proteins.** Shown are Shannon entropy, secondary structure, relative solvent accessibility (RSA) and disordered binding region (DBR) feature scores across five proteins (A) Buc (H0WFA5), (B) Nanos1 (E7FDB3), (C) Nanos3 (Q90WW1), (D) Piwil1 (Q8UVX0), and (E) Tdrd6 (F1R237). Features are normalized on a 0-1 scale, and plotted by averaging along a sliding window of 7 amino acids, except at N- and C-termini where the 4-6 terminal amino acids were used.

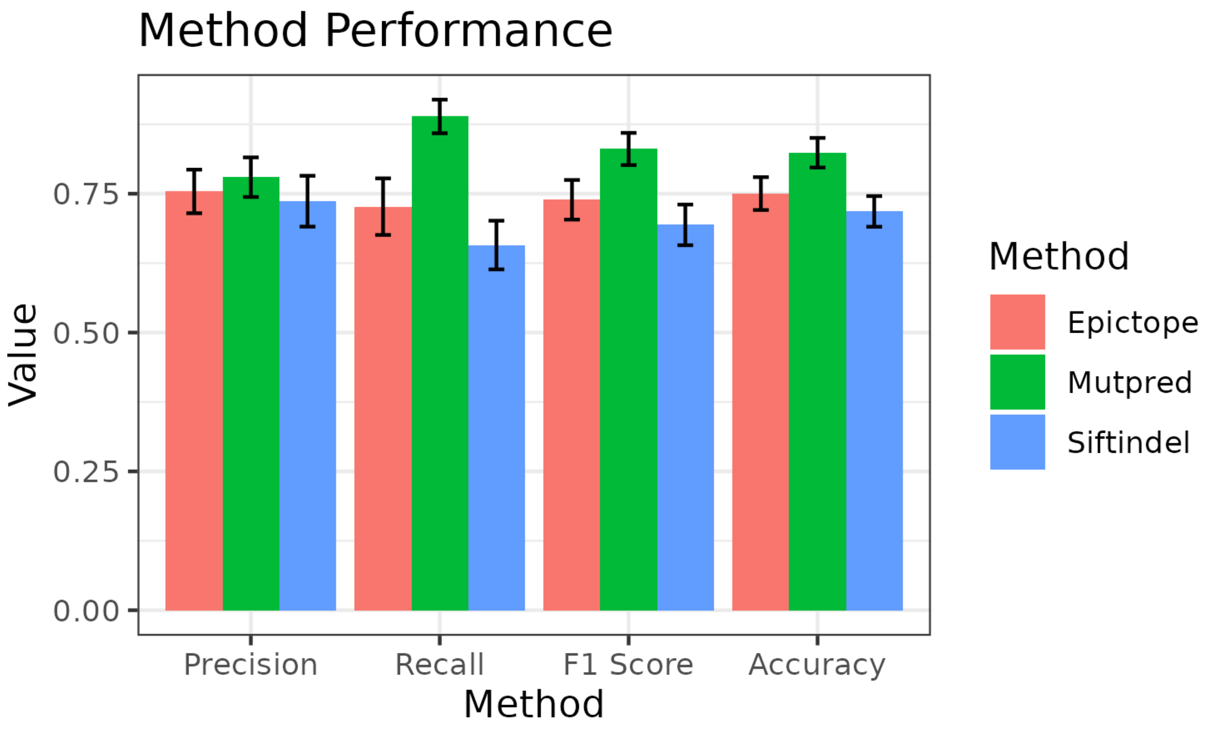


**Figure S6. F1 Score Comparison.** Shown are the precision, recall, F1 and accuracy scores for SIFT-indel, EpicTope, and Mutpred. Error bars indicate standard deviation over 100 replicates of the testing set. SIFT-indel achieved an F1 score of 0.694 (sd = 0.036), EpicTope achieved a score of 0.739 (sd = 0.035), and MutPred2 a score of 0.831 (sd = 0.029).

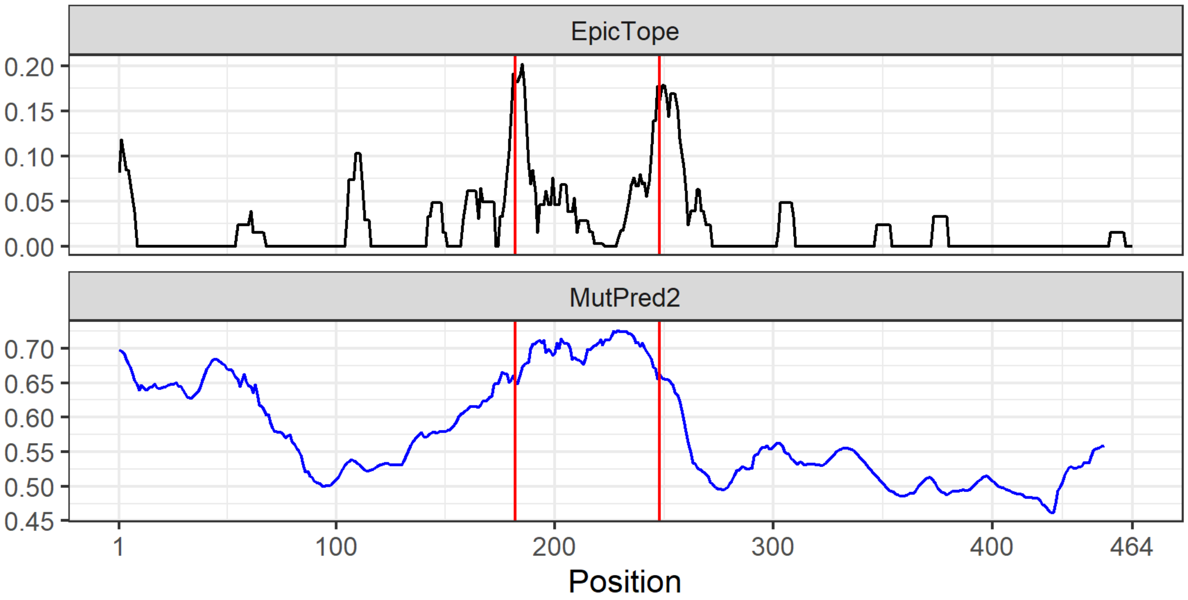


**Figure S7. Predicted Smad5 insertion sites.** Shown are Mutpred2 and EpicTope prediction scores for each position in Smad5. Mutpred2 scores (blue) were generated by subtracting the Mutpred2 pathogenicity score from 1. EpicTope scores are shown in black. Red vertical lines indicate the best-suited EpicTope predicted positions, S181 and S247. Mutpred2 scores are higher for the N-terminus than either internally tagged position. Note that the y-axes scores are not comparable across methods, since MutPred provides a machine learning-based confidence score (0-1), while EpicTope has a score based on its component function as elaborated in the text. EpicTope scores across the protein sequence may vary slightly due to changes in the software comprising the pipeline (IUPred2A and reference database of DSSP). EpicTope and Mutpred2 scores are generated within a moving average window of 7 amino acids. Note that Mutpred2 does not generate scores at C-termini.

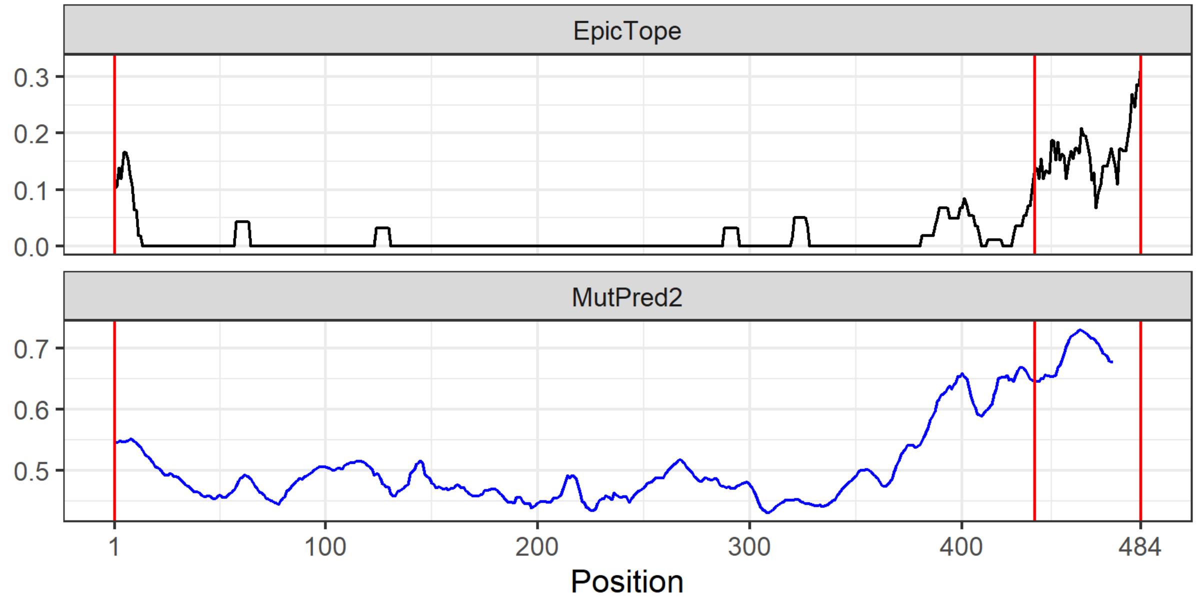


**Figure S8. Predicted Hdac1 tag insertion sites.** Shown are Mutpred2 and EpicTope prediction scores for each position in Hdac1. Mutpred2 scores (blue) were generated by subtracting the Mutpred2 pathogenicity score from 1. EpicTope scores are shown in black. Red vertical lines indicate the three best EpicTope predicted tag positions, A434, as well as the N- and C-termini. EpicTope and Mutpred2 scores are generated within a moving average window of 7 amino acids. Note that Mutpred2 does not generate scores at C-termini.

**Smad5 N-terminal V5 (pBB9)**

```
ATGGGCAAGCCTATCCCAAACCCTCTGCTGGGCCTGGACTCCACAGGTGGTTCTGGTATGACCTCCATG
 M  G  K  P  I  P  N  P  L  L  G  L  D  S  T  G  G  S  G  M  T  S  M
                                                         1
```

**Smad5 S181-V5 (pBB6)**

```
CAGCAGCACAGCGGCAAGCCTATCCCAAACCCTCTGCTGGGCCTGGACTCCACAGGAGGAAGCTCC
 Q  Q  H  S  G  K  P  I  P  N  P  L  L  G  L  D  S  T  G  G  S  S
          181                                          182
```

**Smad5 S247-V7 (pBB8)**

```
GAGACTGGTAGCGGCAAGCCTATCCCAAACCCTCTGCTGGGCCTGGACTCCACAAGCCTGGCTCCT
 E  T  G  S  G  K  P  I  P  N  P  L  L  G  L  D  S  T  S  L  A  P
          247                                          248
```

**Smad5 C-terminal V5 (pBB7)**

```
TCTTCTGTCTCGGGTGGTTCTGGTGGCAAGCCTATCCCAAACCCTCTGCTGGGCCTGGACTCCACATAA
 S  S  V  S  G  G  S  G  G  K  P  I  P  N  P  L  L  G  L  D  S  T  *
          464
```

**Smad5 S181-3xV5 (pKT1)**

```
CAGCAGCACAGCGGCAAGCCTATCCCAAACCCTCTGCTGGGCCTGGACTCCACAGGaAAa
 Q  Q  H  S  G  K  P  I  P  N  P  L  L  G  L  D  S  T  G  K
          181

CCaATtCCcAACCCTCTcCTGGGtCTGGAtagCACcGGaAAGCCaATCCCtAAtCCcCTG
 P  I  P  N  P  L  L  G  L  D  S  T  G  K  P  I  P  N  P  L

CTcGGaCTGGACTCtACtGGAGGAAGCTCC
 L  G  L  D  S  T  G  G  S  S
                  182
```

**Smad5 S247-3xV5 (pKT2)**

```
GAGACTGGTAGCGGCAAGCCTATCCCAAACCCTCTGCTGGGCCTGGACTCCACAGGaAAa
 E  T  G  S  G  K  P  I  P  N  P  L  L  G  L  D  S  T  G  K
          247

CCaATtCCcAACCCTCTcCTGGGtCTGGAtagCACcGGaAAGCCaATCCCtAAtCCcCTG
 P  I  P  N  P  L  L  G  L  D  S  T  G  K  P  I  P  N  P  L

CTcGGaCTGGACTCtACtAGCCTGGCTCCT
 L  G  L  D  S  T  S  L  A  P
                  248
```

**Hdac1 N-terminal V5**

```
ATGGGTAAGCCTATCCCTAACCCTCTCCTCGGTCTCGATTCTACGGGTGGTTCTGGTGCTGAGTTCTCA
 M  G  K  P  I  P  N  P  L  L  G  L  D  S  T  G  G  S  G  A  E  F  S
 1
```

**Hdac1 S434-V5**

```
CGGAGAAACGCAGGTAAGCCTATCCCTAACCCTCTCCTCGGTCTCGATTCTACGGCCAATTACAAG
 R  R  N  A  G  K  P  I  P  N  P  L  L  G  L  D  S  T  A  N  Y  K
          434                                          435
```

**Hdac1 C-terminal V5**

```
TTAAAAACAGTGGGTGGTTCTGGTGGTAAGCCTATCCCTAACCCTCTCCTCGGTCTCGATTCTACGTGA
 L  K  T  V  G  G  S  G  G  K  P  I  P  N  P  L  L  G  L  D  S  T  *
                                                                   484
```

**Figure S9**. Partial sequences of constructs used for rescue experiments. DNA sequences are shown above protein translation in bold. Sequences corresponding to native *smad5* and *hdac1* are highlighted in yellow, GGSG linker is highlighted in grey and V5 is highlighted in aqua. Amino acid numbers are shown in red. Lowercase letters in 3xV5 constructs indicate silent substitutions introduced to make the three tag repeats non-identical to facilitate cloning.

**Supplementary Table 1. Primers used to introduce V5 tags into *smad5* and *hdac1*.** Sequences corresponding to the V5 tag are highlighted in aqua, GGSG linker - in grey, and *smad5* and *hdac1* - in yellow. Lowercase letters denote silent substitutions introduced to make sequences less repetitive.

| Primer | Insertion | Sequence 5'->3' |
|---|---|---|
| V5Smad-F1 | N-terminal V5 | GGCAAGCCTATCCCAAACCCTCTGCTGGGCCTGGACTCCACAGGTGGTTCTGGTATGACCTCCATGTCTAGTCTGTT |
| V5Smad-R1 | N-terminal V5 | ACCAGAACCACCTGTGGAGTCCAGGCCCAGCAGAGGGTTTGGGATAGGCTTGCCCATAGTGCTGGGCTGCACCAGGAAG |
| V5Smad-F2 | S181-V5 | GGCAAGCCTATCCCAAACCCTCTGCTGGGCCTGGACTCCACAGGAGGAAGCTCCTTCCCCATC |
| V5Smad-R2 | S181-V5 | TGTGGAGTCCAGGCCCAGCAGAGGGTTTGGGATAGGCTTGCCGCTGTGCTGCTGGAAGGACTC |
| V5Smad-F3 | S247-V5 | GGCAAGCCTATCCCAAACCCTCTGCTGGGCCTGGACTCCACAAGCCTGGCTCCTCAGAACATG |
| V5Smad-R3 | S247-V5 | TGTGGAGTCCAGGCCCAGCAGAGGGTTTGGGATAGGCTTGCCGCTACCAGTCTCCATGGACTGA |
| V5Smad-F4 | C-terminal V5 | GGTGGTTCTGGTGGCAAGCCTATCCCAAACCCTCTGCTGGGCCTGGACTCCACATAATGATGGGCTGACCTGGGAGG |
| V5Smad-R4 | C-terminal V5 | TGTGGAGTCCAGGCCCAGCAGAGGGTTTGGGATAGGCTTGCCACCAGAACCACCCGAGACAGAAGAGATGGGGTTC |
| Smad5S181-3xV5-F1 | S181-3xV5 | ATtCCcAACCCTCTcCTGGGtCTGGAtagCACcGGaAAGCCaATCCCtAAtCCcCTGCTcGGaCTGGACTCtACtGGAGGAAGCTCCTTCCCCATC |
| Smad5S181-3xV5-R1 | S181-3xV5 | CAGaCCCAGgAGAGGGTTgGGaATtGGtTTtCCTGTGGAGTCCAGGCCCAGCAGAGGGTTTGGGATAGGCTTGCCGCTGTGCTGCTGGAAGGACTC |
| Smad5S247-3xV5-F1 | S247-3xV5 | ATtCCcAACCCTCTcCTGGGtCTGGAtagCACcGGaAAGCCaATCCCtAAtCCcCTGCTcGGaCTGGACTCtACtAGCCTGGCTCCTCAGAACATG |
| Smad5S247-3xV5-R1 | S247-3xV5 | CAGaCCCAGgAGAGGGTTgGGaATtGGtTTtCCTGTGGAGTCCAGGCCCAGCAGAGGGTTTGGGATAGGCTTGCCGCTACCAGTCTCCATGGACTG |
| Hdac1-F | Hdac1 ORF | CGACGCCACCATGGCGCTGAGTTCTCAAG |
| Hdac1-R | Hdac1 ORF | AGATCCGCCGTCACACTGTTTTTAATTCTTCTTTTGG |
| V5Hdac1-N-F | N-terminal V5 | CCCTAACCCTCTCCTCGGTCTCGATTCTACGGGTGGTTCTGGTGCGCTGAGTTCTCAAGGAACAAAGAAGAAAGTTTGC |
| V5Hdac1-N-R | N-terminal V5 | CCGAGGAGAGGGTTAGGGATAGGCTTACCCATGGTGGCGTCGAGTTAGATCTGCCAAAGTTGAGCG |
| V5Hdac1-434-F | A434-V5 | CTAACCCTCTCCTCGGTCTCGATTCTACGGCCAATTACAAGAAGCCAAAACGAGTG |
| V5Hdac1-434-R | A434-V5 | GAGGAGAGGGTTAGGGATAGGCTTACCTGCGTTTCTCCGTCCGCCCTGGC |
| V5Hdac1-C-F | C-terminal V5 | GGTAAGCCTATCCCTAACCCTCTCCTCGGTCTCGATTCTACGTGACCGCGGATCTGGTTACCACTAAAC |
| V5Hdac1-C-F | C-terminal V5 | GAGGAGAGGGTTAGGGATAGGCTTACCACCAGAACCACCCACTGTTTTTAATTCTTCTTTTGGCCCCTTTGTGTCCATTTTCTCTTC |
| pT3TS-F | pT3TS Backbone | CACCAGTCACAGAAAAGCATCTTACGGATGGCATGACAGTAAGAG |
| pT3TS-R | pT3TS Backbone | CGTAAGATGCTTTTCTGTGACTGGTGAGTACTC |
